# Mixed Selectivity of Subthalamic Nucleus Neurons in Encoding Motor and Reward Behaviors

**DOI:** 10.1101/2025.02.12.637797

**Authors:** Bing-Shiuan Wu, Min-Yuan Ming, Yu-Wei Wu

## Abstract

The subthalamic nucleus (STN) plays a critical role in modulating motor and cognitive functions within the basal ganglia, with its dysfunction being implicated in movement disorders such as Parkinson’s disease. However, the behavioral representations of individual STN neurons remain incompletely understood. Using *in vivo* two-photon calcium imaging in behaving mice, we systematically mapped the activity of single STN neurons across diverse behavioral contexts, including locomotion, licking, and reward-driven actions. Our findings reveal that STN neurons exhibit mixed selectivity, encoding multiple behaviors with distinct temporal dynamics and excitatory or inhibitory response patterns. This mixed selectivity allows the STN to robustly encode motor parameters such as locomotion speed and licking intensity while integrating contextual information from different behavioral states. Comparisons with the adjacent zona incerta (ZI) revealed distinct encoding properties: while both regions represent locomotion, STN neurons more faithfully track motor states, whereas ZI neurons exhibit prolonged calcium events with weaker movement correlations. Population-level analysis showed STN activity in a low-dimensional neural manifold, with components linked to movement velocity and licking intensity. Notably, locomotion encoding in STN was context-dependent, diverging when movements were internally generated versus reward-modulated. Together, these findings highlight the specialized yet flexible role of the STN in integrating motor and reward-related signals, supporting a framework in which STN neurons contribute to motor control through multiplexed and context-dependent encoding. This work provides new insights into the functional organization of basal ganglia circuits and has implications for understanding STN’s role in both physiological and pathological conditions.

## Introduction

The cortico-basal ganglia-thalamic loops have been described to parallelly transduce signals from varied functional domains, including those of sensorimotor, associative, and limbic systems ^1-3^. Within the loops, while most of the basal ganglia sub-structures send inhibitory outputs to their downstream targets, the subthalamic nucleus (STN) is the only excitatory component ^4^. In the pathological condition, it has long been observed that the STN shows pathological hyperactivity in Parkinson’s disease (PD). In PD patients, deep-brain stimulation (DBS) targeting STN can effectively alleviate motor symptoms ^5,6^. However, the STN neuronal population is heterogeneous in connectivity and gene expression ^7^. How the diverse STN neurons are manipulated by the DBS to reach the therapeutic effect, and how these neurons cooperatively representing behavioral states remain controversial.

The function of STN is traditionally considered as motion suppression, which is supported by many manipulating and recording experiments ^8-12^. The stop signal is suggested to be especially conveyed via the frontal cortico-STN connection ^12,13^. However, in addition to stop responses, other studies have identified moving-correlated activity in STN through *in vivo* recordings in monkeys and mice ^14,15^. Also, transient opto-stimulation of STN is sufficient to generate short-latency body movements, which are linearly scaled with the stimulus frequency ^16^. These suggest that apart from motion suppression, the STN neurons also participate in pro-movement generation.

Several pieces of *in vivo* recording works in primates suggest the multifunctionality in the STN. It has been identified in humans that some STN units elevate firing on the moving onset, while others respond to the stop cue ^17^. Further, “movement cells”, “stop cells”, “switch cells”, and even “torque cells” are recognized in monkeys’ STN by performing Go/No-Go arm movement tasks ^18^. Apart from the motor functions, non-motor roles of the STN are also identified ^19-22^. Especially, reward amplitude is shown to be encoded by STN neurons ^23,24^, suggesting the limbic function in the STN.

In this study, we hypothesized that STN neurons exhibit multimodal representations that collectively encode motor and reward-related behaviors across different contexts. Using *in vivo* two-photon calcium imaging in head-fixed behaving mice, we confirmed that individual STN neurons display mixed selectivity, encoding multiple behaviors with unique temporal dynamics and excitatory or inhibitory responses. This mixed selectivity allows STN neurons to robustly represent motor parameters, such as locomotion speed and licking intensity, while contextualizing these behaviors with distinct neural activity patterns. Furthermore, we found that STN neurons reliably encode motor states with contextual specificity, differing from the nearby ZI, where neurons demonstrate more varied calcium response patterns and a weaker correlation with movement parameters. These findings highlight the specialized yet flexible role of the STN in regulating motor and reward-related behaviors, contributing significantly to basal ganglia circuit function.

## Results

### *In vivo* single-cell calcium imaging of STN neurons in head-fixed, behaving mice

To achieve single-cell resolution in STN activity recording *in vivo*, a GRIN lens or guide cannula was implanted to target the STN following infection with AAV9-CAG-GCaMP6f infection (Fig. 1a and b; Fig. S1a and S1b). During recordings, the mice were trained to remain head-fixed on a running wheel and allowed to move voluntarily. To enhance behavioral diversity, a 4% sucrose water reward was randomly provided without any additional cues. Calcium activity was measured using two-photon microscopy, and data on wheel rotation, reward timing, and licking events were collected simultaneously in this unstructured task setup (Fig. 1c, d and e). We verified that the implanted mice exhibited normal locomotion in an open-field test, indicating no impairment from the GRIN lens and cannula surgeries (Fig. S1c, S1d). Neuronal activity could be reliably recorded over several months without notable photobleaching or cell death (Fig. S1e), allowing us to accumulate data across multiple sessions.

**Figure 1.**
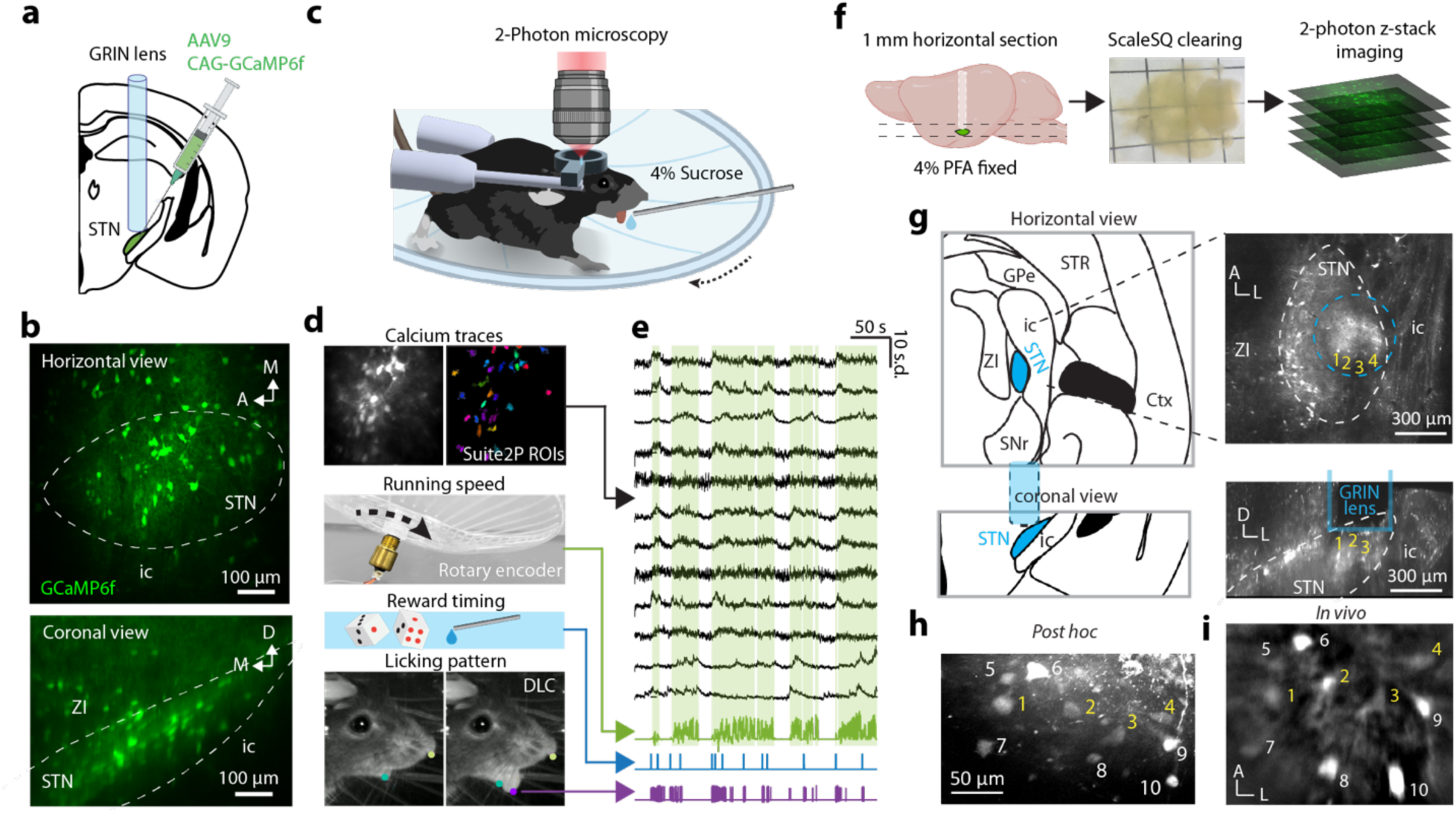
*In vivo* calcium imaging in behaving mice and *post hoc* validation of recorded fields in the STN. **a,** GCaMP6f was expressed in the STN via viral injection, followed by the implantation of a GRIN lens or guided cannula (see Methods for details). **b,** Representative images showing GCaMP6f expression in the STN from horizontal (top) and coronal (bottom) views. **c,** Schematic of the unstructured behavioral task during recording: head-fixed mice ran voluntarily on a running wheel, with a 4% sucrose reward delivered at random intervals. **d**, Calcium traces of individual neurons were extracted using Suite2P; locomotion data were recorded by a rotary encoder, and licking events were detected by DeepLabCut from infrared camera footage. **e,** Example traces of calcium activity aligned with running speed, reward timing, and licking patterns. Green shading indicates movement episodes. **f,** Workflow for *post hoc* validation of recorded fields. **g**-**i,** Represented example for *post hoc* identifying and localizing the *in vivo* recorded cells in the fixed slice. **g**, The GRIN lens position and the anatomical structure of the STN in the fixed brain slice. **h,** The GCaMP6f-expressing neurons in the fixed slice that had been recorded *in vivo*. **i**, Representative averaged image from *in vivo* calcium imaging through GRIN lens. The annotated numbers indicate the same neurons identified in (**h**) and (**i**).

To confirm the recorded regions, mice were perfused, and thick brain slices were sectioned horizontally. Tissue clearing and two-photon z-stack imaging were applied to map the *in vivo* imaged fields onto fixed slices (Fig. 1f). By identifying cells imaged *in vivo* within the slices, we confirmed the anatomical positions of the fields of view (FOVs) (Fig. 1g-j). This *post hoc* validation after multiple recording sessions ensured the accurate capture of single-cell activity in the STN (n = 80 neurons in 4 FOVs) and the surrounding zona incerta (n = 150 neurons in 5 FOVs).

### Majority STN neurons increase activity during locomotio*n*

Locomotion-related increased activity in STN has been documented ^15,25^; however, the detailed temporal patterns and the single-cell responses have not yet been thoroughly analyzed. Further, how the locomotion-responding neurons react to other behaviors remains unexplored. Here we extracted the time points of the initiation and the termination of robust moving episodes (lasting more than 4 seconds), in order to get the “go” (n = 64 responsive cells) and “stop” (n = 72 responsive cells) responses of each cell. The majority of recorded STN neurons are activated during the “go” phase and decrease the activity during the “stop” phase (Fig. 2c and i). However, nuances of calcium responses are observed across cells. When focusing on the single-trial traces, the calcium activity changes across moving bouts appear to be considerably reliable (Fig. 2a, b, g, h, Fig. S2a-f). The trial-averaged activities from the neurons are then subjected to clustering analysis. The features in the trial-averaged calcium activities are extracted in the first few principal components (PCs), which are utilized in hierarchical clustering (Fig. S2g-j).

**Figure 2.**
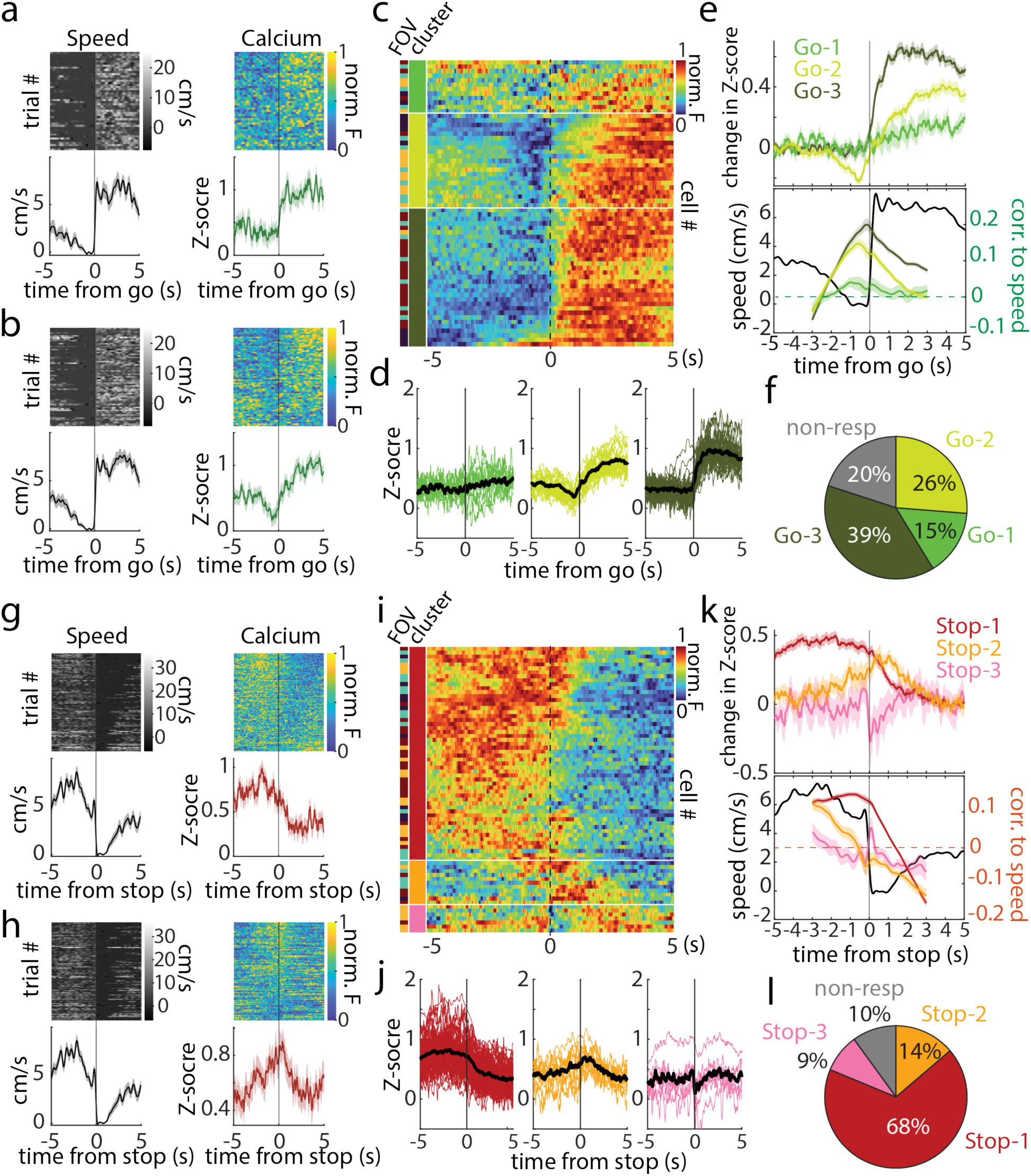
The calcium activities of the STN neurons in response to locomotion initiation and termination are clustered into subtypes. **a**, **b** Two examples of single-trial calcium response aligned to the "go" time points. The calcium activities displayed in the heat map are normalized to a range of 0 to 1 and match the corresponding behaviors. **C** The trial average for each recorded cell is normalized to a range of 0 to 1. Each row is color-coded according to the cell’s corresponding subtype and the field of vision (FOV) it belongs to. **d** Single-cell trial-averaged traces from each cluster are displayed as colored thin lines, while the cluster-averaged traces are represented by black thick lines. **e** The upper panel shows the superposition of the offset cluster-averaged traces, and the lower panel displays the trial-by-trial time-lagging cross-correlation between calcium activity and the locomotion speed. The shaded area in the line plots indicates the SEM. **f** The percentage of each cluster of response types. The gray color indicates the proportion of non-responsive cells. **g**-**l** Same as a-f but for activity aligned to “stop” time points.

Firstly, the cells are clustered into three subtypes according to their “go” responses, with two of them (“Go-2” and “Go-3”) showing robust increased activity (Fig. 2c-e). Detailly speaking, compared with the responses in the “Go-3” subtype, those in the “Go-2” subtype exhibit a brief inhibition before the running onset and a slower ramp-up phase afterward (Fig. 2e, upper panel). These features can also be recognized in the trial-by-trial time-lagging cross-correlation between the calcium traces and the speed, which highlights their temporal relations. The cross-correlation reaches the peak at around zero lags for the “Go-3” subtype. In comparison, the correlation for the “Go-2” subtype peaks at negative time lags and drops steeper when the calcium traces are shifted with positive time lags, which reflect the slow ramping and the brief inhibition of the activity, respectively (Fig. 2e, lower panel).

Secondly, in the cellular subtypes defined by the “stop” responses, a majority of the cells (“Stop-1”) show a similar pattern of decaying activity in response to the dropping speed (Fig. 2i and j). The activities from this group of cells display the highest correlation to the speed change. A smaller subgroup (“Stop-2”) exhibits a transient activation around stopping, which is inversely correlated to the speed. The third cluster (“Stop-3”) shows a short activity drop immediately after stopping, which gives rise to a peak in the correlation (Fig. 2k). However, these activities are relatively weaker and noisier than others (Fig. 2j). In summary, in response to transitions in the locomotion states, the activities of most STN neurons align with changes in speed. Nevertheless, nuances of the temporal patterns can be recognized among different sub-populations within the STN.

### STN neurons display diverse types of activity in response to licking and random reward

In addition to locomotion, licking behavior was monitored. Licking intensity, reflecting the frequency, is quantified by the kernel-convolution of each licking event (Fig. 3b). Licking trials were categorized as reward-driven or spontaneous ones, based on the presence or absence of reward delivery, respectively (Fig. 3a, c, and d). Almost every reward delivery was followed by burst-like, continuous licking, while single, sparse licks occurred spontaneously without reward (Fig. 3c and d, licking traces). The calcium responses in a single neuron can vary depending on the type of licking behavior. For instance, whereas an STN neuron might activate at sparse spontaneous licking (Fig. 3c and d, upper parts), the same cell may display a strong, prolonged activation or an inhibition in response to reward-driven licking (Fig. 3c and d, lower parts). This indicates that the calcium activity during burst-like reward-driven licking is not linearly scaled from that during a single lick without reward. Furthermore, the combination of responses to licking and locomotion varied across cells (Fig. S3a-h).

**Figure 3.**
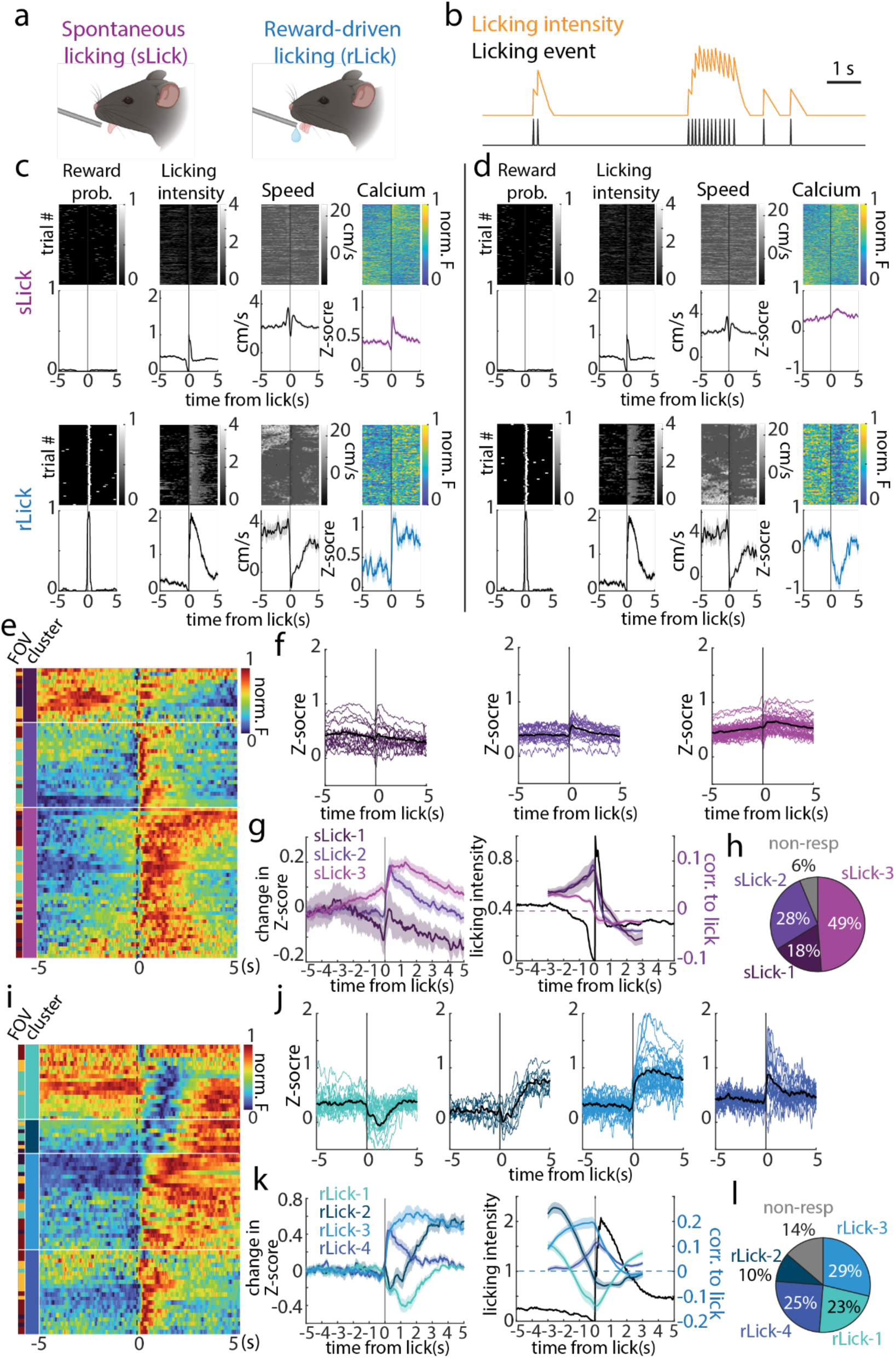
The calcium responses to licking onsets are not interdependent between spontaneous and reward-driven licking, which are clustered into four unique subtypes. **a** Licking onsets can be separated into spontaneous and reward-driven ones. **b** “Licking intensity” is the convoluted traces of single licking events with a linearly-decaying kernel. **c**, **d** Single neuron examples of the behavior changes and the calcium responses under onsets of spontaneous and reward-driven licking. The upper panels are the licking onsets without reward (sLick), while the lower panels are the trials with reward (rLick). The traces are aligned to the licking onsets, with licking intensity patterns and locomotion speed being shown with the corresponding calcium activity. The shaded area in the line plots indicates the SEM. **e-h** Same as fig. 2c-f but for spontaneous licking. **i-l** Same as fig. 2c-f but for reward-driven licking.

Through the same clustering approach (Fig. S3i-l), responses to both types of licking can also be grouped into subtypes. Spontaneous licking (sLick), like self-paced locomotion, is considered as an internally driven behavior. This type of licking is usually accompanied by a modest elevation in calcium activity in most STN neurons (n = 75 responsive neurons). It reveals that some of the neurons exhibit faster decay (sLick-2) while others are more prolonged (sLick-3) (Fig. 3e-f). For reward-driven licking (rLick), on the other hand, the calcium responses can be classified into four subtypes (n = 69 responsive neurons), each with unique dynamic patterns (Fig. 3i-l). Specifically, a subtype of STN neurons shows an inhibition in the activity upon licking onsets (rLick-1). Other three types exhibit activation with distinct temporal dynamics, which can be described as delayed (rLick-2), prolonged (rLick-3), and transient (rLick-4) activation, respectively (Fig. 3k). Notably, although rLick is accompanied with locomotion stop if the reward was given during running, the responses are not dependent to the moving speed when observing the single trials (Fig. 3c and d, lower panels), suggesting the activities are not representing deceleration *per se*. It is worth noting that the diversity of response types is observed within a single recording field (Fig. 3i), indicating that the subtypes are spatially intermingled.

### Cross-talking of response subtypes defined by different behaviors indicates multiple representations in single STN neurons

In the recording sessions, each cell was exposed to the four behavioral conditions: go, stop, reward-driven licking, and spontaneous licking. Next, we aim to explore how the subtypes defined by different behaviors interact at single-cell level. In other words, how the activity of an individual cell responds to these various behaviors is going to be investigated.

To begin with, the activity traces of each cell is projected onto the two-dimensional subspace defined by the first and second principal components (PCs) of the “rLick” response traces—which show more unique patterns. In this subspace, “rLick” response traces with similar features are positioned closer together than those with distinct waveforms (Fig. 4a, upper-left panel). With the same coordinates in the subspace, each representing a single cell, the calcium traces for other behaviors are plotted (Fig. 4a, other three panels). Each cell’s trace is colored based on its belonging subtype, while the gray traces represent cells that do not show a significant response to the behavior condition.

**Figure 4.**
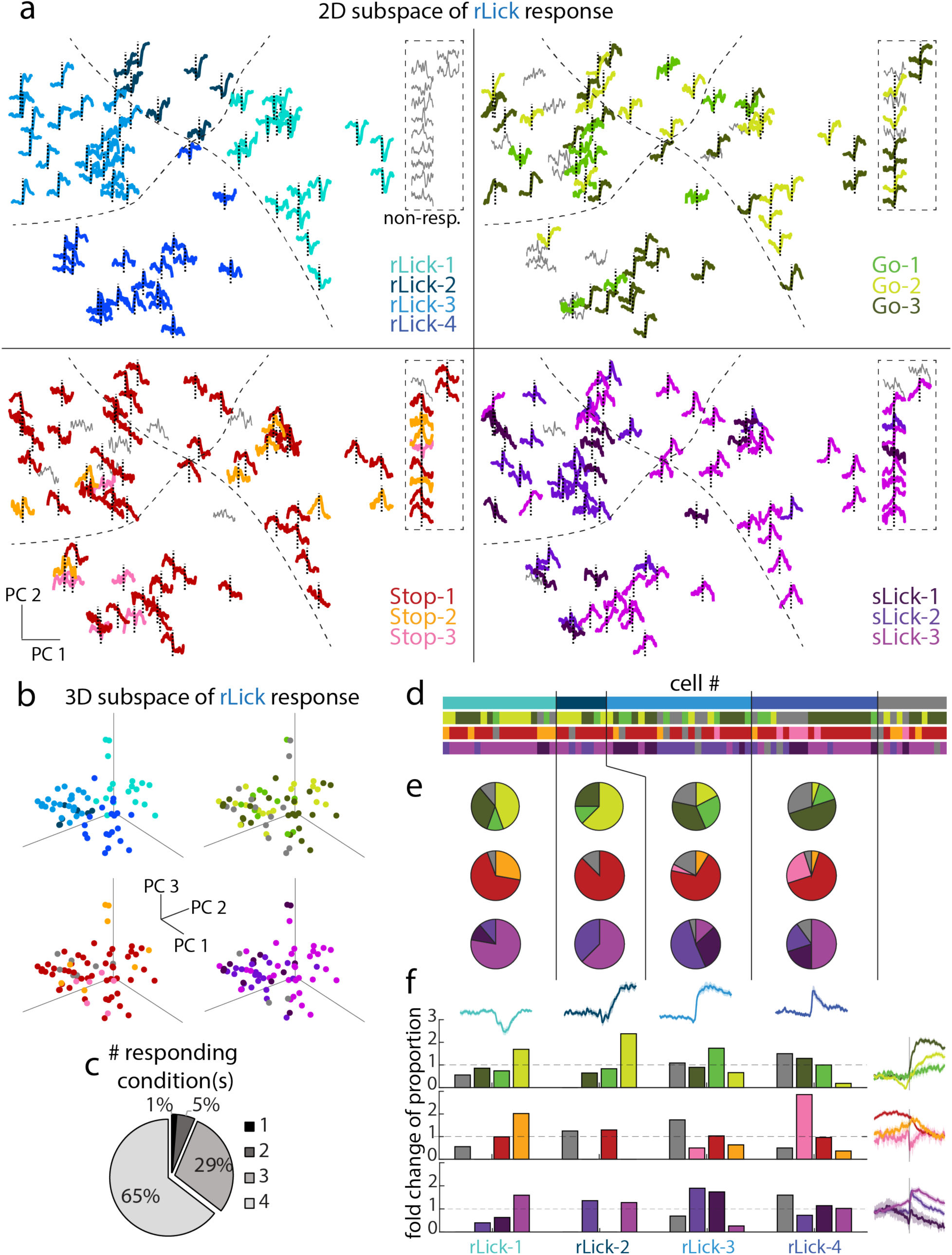
Cross-talking of single-cell calcium response clusters across behavioral conditions. **a** Mapping of calcium response of each cell across behaviors in the “reward-driven licking” response subspace. The traces in the four panels are the responses to the corresponding behaviors, and the positions are defined by the PC 1 and PC 2 of their “reward-driven licking” response. The gray traces represent the non-significant responses and are placed in the upper-right box. The traces are color-coded according to their subtypes. **b** The single-cell distribution in the 3D subspace defined by the PC 1, PC 2, and PC 3 of their “reward-driven licking” response. The dots are color-coded according to their subtypes. **c** The pie chart representing the proportions of cells that are significantly responsive in one to four behavioral conditions. **d** Single-cell cluster distributions. In the four horizontal bars, the colors in each column represents the clusters to which a cell belongs. The order of the cells is sorted according to the subtypes 1-4 of the “rLick” response. The gray color represents the non-significant response. N=80 cells. **e** Proportional distribution of different response patterns across behaviors in each “rLick” subtype. **f** The fold-change of the subtype proportion within a specific "rLick" cluster (e) compared with its overall distribution in all recorded cells (fig. 2f, 2l, 3h, and 3l). A fold-change value of 1 indicates that the proportion of the subtype in the specific cluster is the same as its overall proportion.

This visualization clearly demonstrates that the response subtypes for other behaviors are not segregated in the “rLick” response subspace, indicating that cells with similar “rLick” responses can exhibit divergent activities under other behaviors. Moreover, some STN cells are not statistically responsive (gray traces) under “rLick” but respond to other behaviors. These intermingled distributions are also seen when projecting the responses onto the subspace of “Go” response (Fig. S4a), and that with higher dimensions, which taking more detailed features into consideration (Fig. 4b and S4b). Under the four behavioral conditions, a majority of cells are significantly responsive and fall into one of those subtypes (Fig. 4d). As much as 94% of the recorded cells show responses to more than three behavioral conditions, with 65% being responsive to all of them (Fig. 4c). This suggests that most of the STN neurons display multiple behavioral representations.

To determine to what extent the subtypes defined by the responses to reward-driven licking correlate with categorizations under other conditions, the proportion of clusters for other behaviors within each “rLick” category is quantified (Fig. 4d, e and f). Two trends of correspondence among conditions can be summarized. Firstly, cells belonging to the inhibition type in reward response (rLick-1) display a higher percentage of cells that show inhibited activity before “go” (Go-1). This type also contains more “Stop-2” cells (transient activated at stopping) and “sLick-3” cells (slow-decaying activation at spontaneous lick) (Fig. 4d, e and f). Secondly, cells with rapid rising in response to reward (rLick-3 and rLick-4) include a higher proportion of cells with weak and noisy (e.g. Go-1, Stop-3, and sLick-1), or even non-significant, responses under other behavioral conditions (Fig. 4d, e and f; also shown in the dashed line box of Fig. 4a, upper-right panel).

These results suggest that a single STN neuron can represent multiple behavior conditions. While the representations are partially interdependent among behaviors, exceptions that deviate from this pattern also cannot be ignored. In brief, cells with similar responses under one behavior can act differently under other conditions. That is to say, simple classifications are inadequate to fully explain the diverse representations observed in all the recorded STN neurons.

### Population dynamics reveal essential components of the diverse representations in the STN neurons

Regarding the mixed selectivity of the multiple representations across the STN neurons, single-cell clustering falls short of elucidating the underlying computational principles. Therefore, we seek to explore the functional population dynamics of the STN. Neuronal activities can be described in a high-dimensional neural space, where each axis corresponds to the activity of an individual cell. The individual cells do not fire independently during the computation process, but rather with a certain coordination. Accordingly, activities within this high-dimensional space tend to occupy a restricted, low-dimensional subspace ^26^ (Fig. 5a), with each axis being the weighted combination of all neuronal activities (Fig. 5b and Fig. S5). We performed dimensionality reduction of the neuronal activities in order to visualize the temporal evolution of the population dynamics under each condition.

**Figure 5.**
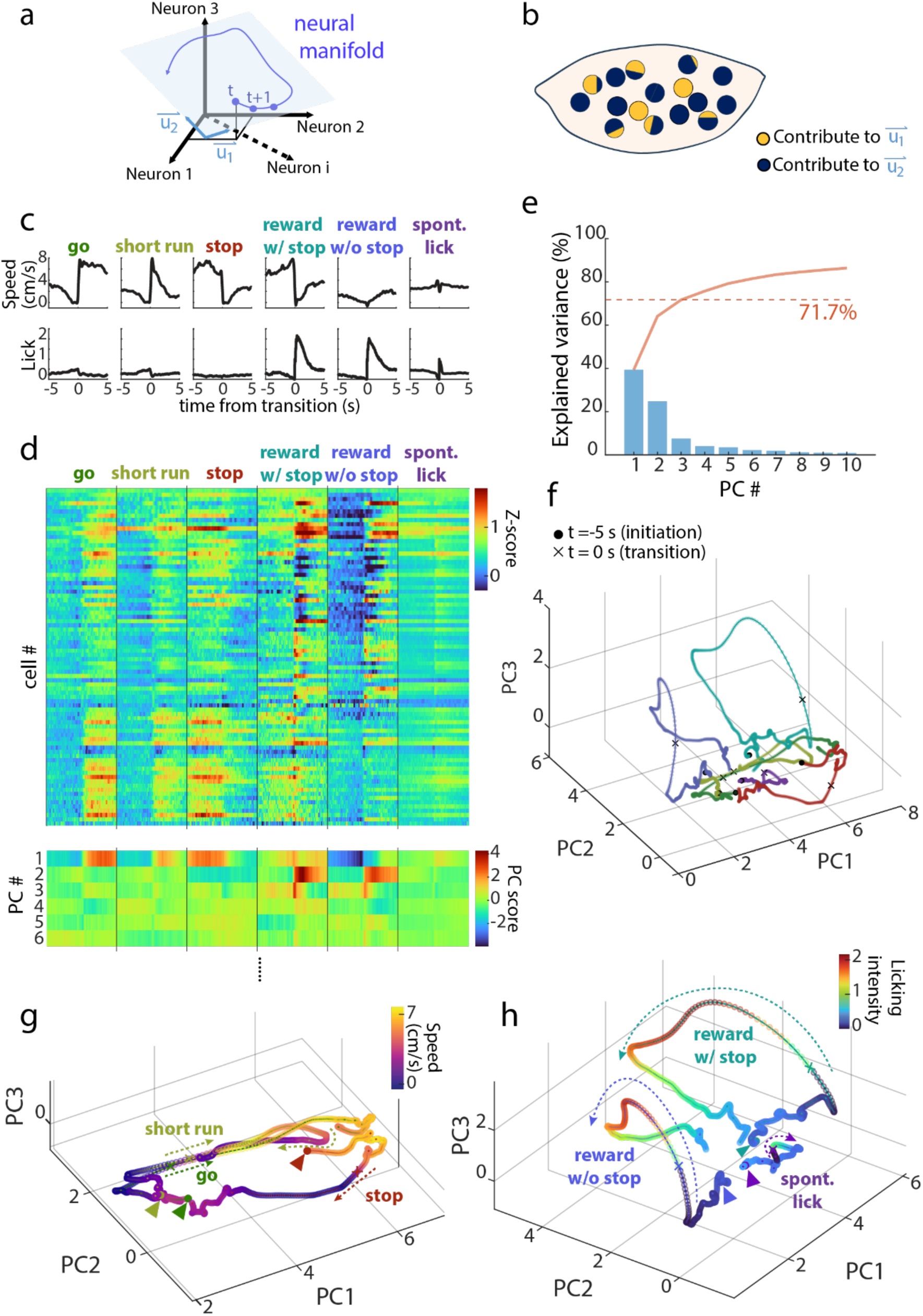
Population dynamics of STN neurons under various behavioral conditions. **a**, **b** Schematic of the concept of neural subspace in the dynamical system. **c** Behavioral parameters, including trial-averaged locomotion speed and licking intensity, under the six analyzed behavioral conditions. **d** Trial average of z-scored calcium activity of all cells under all six behavioral conditions (upper panel) was transformed in to PC scores along time points. Dynamics of the first 6 PCs is shown (lower panel). Only the first 3 PCs (71% variance) are used for the manifold. **e** The proportion of variation being explained in each principal component. The orange line represents the cumulative distribution. **f** Neural trajectories under the six behavioral conditions in the three-dimensional subspace established by PC 1, 2, and 3. **g** The neural trajectories for the three locomotion conditions. The trajectories are color-coded based on the corresponding averaged speed. **h** The trajectories for the three licking conditions. The colors represent the licking intensity.

The trial-averaged calcium activities under different behavioral conditions, including those previously mentioned and the “short run” trials where moving bouts last less than 4 seconds, are subject to analysis. Further, the “reward-driven licking” events are separated into those with robust stopping from running and those without high initial speed (Fig. 5c, “Reward with stop” and “Reward without stop”). Through PCA, the high-dimensional single-cell activities (n = 80) at any given time point are transformed into principal components (Fig. 5d). The first 3 PCs collectively explain more than 70% of the variance (Fig. 5e), allowing the time-evolving trajectory of activity under the five behaviors to be visualized in a three-dimensional subspace (Fig. 5f).

In the behaviors “go”, “short run”, and “stop”, the neural trajectories majorly travel along the PC 1 axis, with some deviations in PC 2. Specifically, during the “go” behavior, the trajectory moves from lower to higher values on the PC 1 axis, whereas during the “stop” behavior, it moves in the opposite direction. Notably, the trajectory for the “short run” initially resembles that of the “go” behavior but reverses direction as the speed decreases (Fig. 5g).

On the other hand, immediately following the onset of “reward-driven licking”, the trajectory extends largely along PC 2 and PC 3. In consistent with locomotion modulation mentioned above, prior to licking onsets and reward delivery, the trajectory initiates with a higher PC1 value when the reward is delivered while moving (“Reward with stop”), compared to when the reward is given while stationary (“Reward without stop”). In addition, during “spontaneous licking”, the trajectory forms a small loop on the PC 1-2 plane (Fig. 5h). In brief, the neural dynamics in response to reward, which involves an external information input, diverge from the dynamics during locomotion in the subspace.

To examine the relationship between behavioral parameters and the principal components, the trajectories are color-coded based on the average locomotion speed (Fig. 5g and 6a) or licking intensity (Fig. 5h and 6g) at the corresponding time points. There is a tendency for higher speeds to correspond with higher values on PC 1 (Fig. 6a), which shows the strongest correlation compared to other principal components (Fig. 6d and f). On the other hand, the licking intensity displays the highest positive correlation to PC 2 values (Fig. 6g and h).

**Figure 6.**
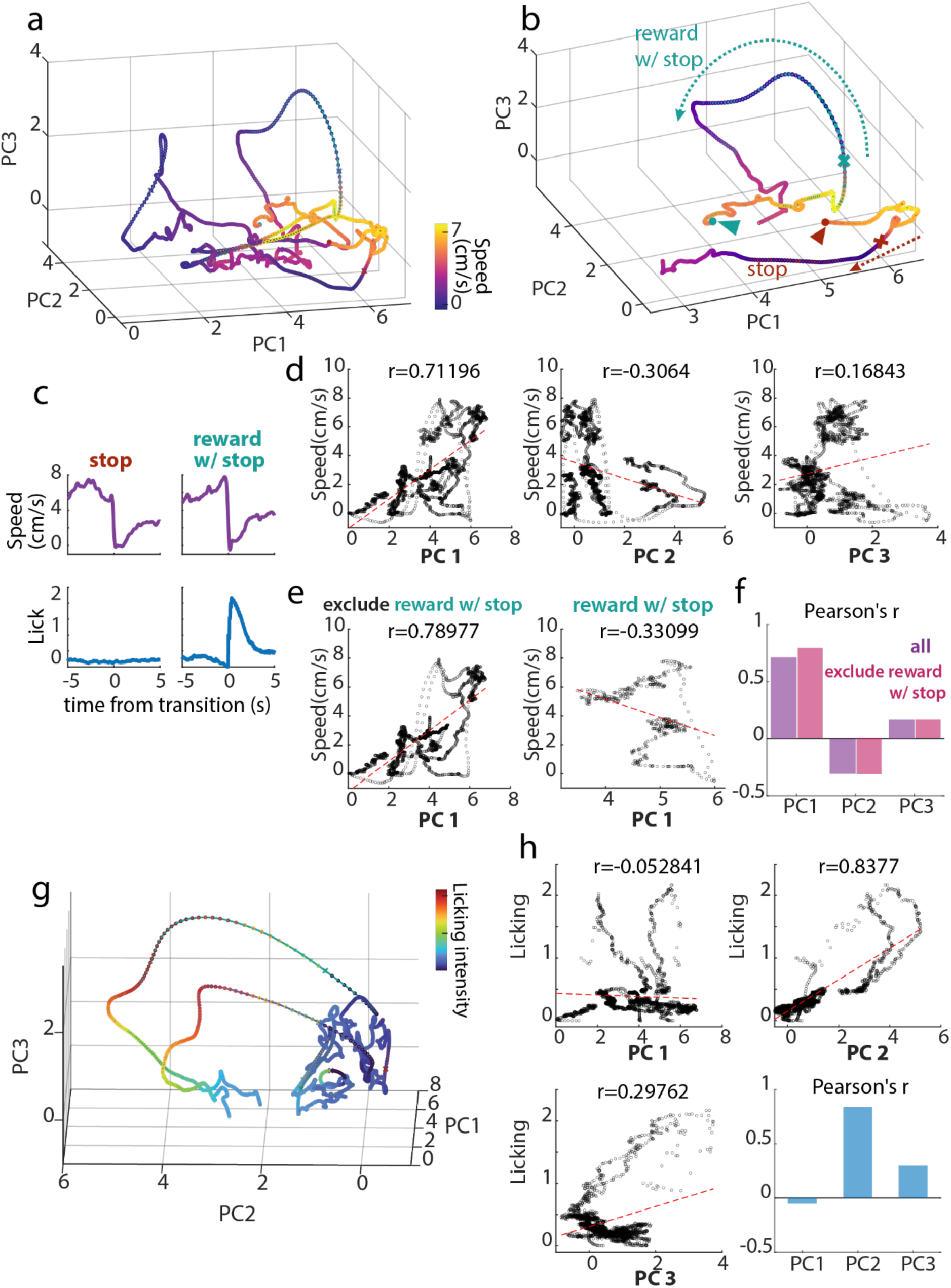
Principle components are linked to behavioral parameters in a context-dependent manner. **a** Neural trajectories for the activities across all six behavioral conditions depicted in Fig. 5, which are colored by locomotion speed. **b** The trajectories for “stop” and “reward with stop” conditions are colored by the speed, showing that similar speed changes can have divergent PC 1 changes. **c** “Stop” and “reward with stop” conditions share similar averaged speed patterns. **d** Values of each principal component are plotted against the corresponding speed. The red dashed lines are the fitted regression line. The r values are the Pearson’s correlation coefficients between PCs and speed. **e**, **f** PC 1 and speed are more correlated when the “reward with stop” condition is excluded, indicating the speed encoding is weak specifically under this context. **g** Neural trajectories for the activities across all six behavioral conditions, which are colored by licking intensity. **h** Values of each principal component are plotted against the corresponding licking intensity. The lower-right panels present Pearson’s correlation coefficients to quantify the relationships between the principal components and licking.

The speed-PC 1 correlation, however, demonstrates context-dependency. Both self-paced “Stop” and “Reward with stop” conditions show similar locomotion deceleration, but one is spontaneous and the other is reward-induced and accompanied by reward-driven licking (Fig. 6c). While the self-paced “Stop” reduces the moving speed along with the decrease of PC 1 value, the neural dynamics during the deceleration of “Reward with stop” behavior travel along PC 2 and PC 3 instead of the PC 1 axis (Fig. 6b). Indeed, while the dynamics of the conditions display higher correlation with PC 1 when excluding the “Reward with stop” trajectory (Fig. e and f). This indicates that the external reward signal intervenes the basic representation of PC 1 in locomotion speed.

The results demonstrate that within the population, one component (PC 1), which reflects a specific weighted combination of single-cell activities, encodes the movement velocity. Importantly, this representation is shown to be context-dependent, with the neural dynamics under the reward condition may deviate from the tendency. Indeed, single STN cells can react distinctively during locomotion deceleration depending on whether a reward is present or not (Fig. S6). Meanwhile, another component (PC 2) encodes licking intensity. There is remaining activity variance in PC 3, the physiological significance of which remains unclear based on current evidence. However, we hypothesize that this dimension may encode information related to either reward prediction or action state transition, as the trajectory for "reward-driven licking," compared to other conditions, travels more extensively along this axis (Fig. 5f and h). In summary, STN neurons, instead of forming functional clusters across conditions, coordinate and emerge certain hidden factors to compute and represent various motor behaviors.

### STN neurons represent behavioral conditions distinctively compared to the zona incerta

To address whether the representations we described are unique in the STN, we analyzed the responses from neurons located outside, but adjacent to, the STN. Through post hoc histology, some recording fields are considered not in the STN and been excluded from the aforementioned analysis (Fig. S7a-d). These neurons are located medial or dorsal to the STN, where falls in to the region of zona incerta (ZI) (Fig. 7a, b, and S7d). We found that the calcium activity in this region is more diverse than those within the STN, containing cells exhibiting long calcium events, with some of them being movement irrelevant (Fig. 7b).

**Figure 7.**
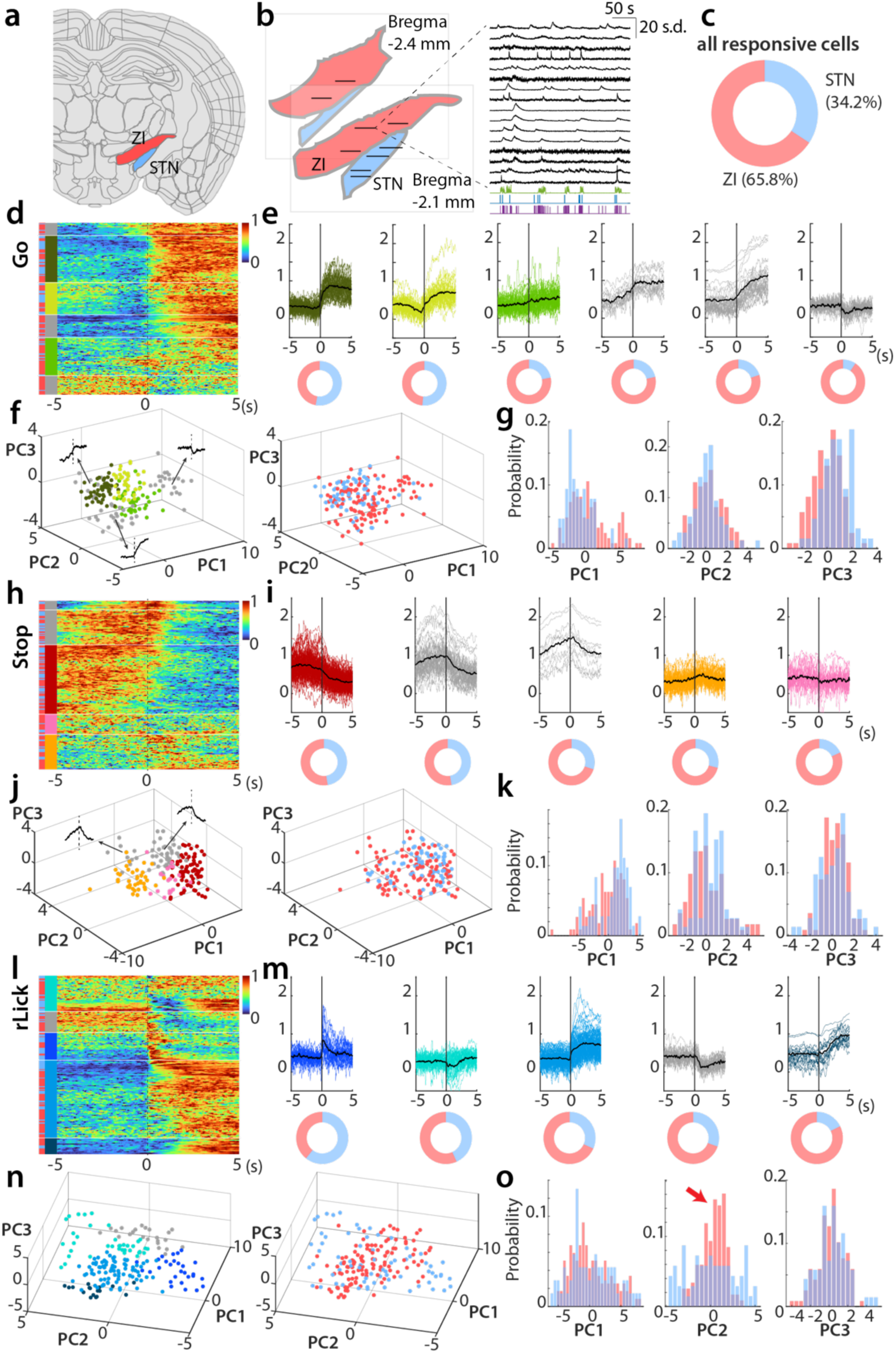
Zona incerta neurons represent behavioral contexts differently from STN. **a, b** Recording fields include both STN and ZI regions. Each horizontal line represents a recorded plane. The right panel in (b) demonstrates the example traces from ZI. **c** Proportional distribution of the number of responsive cells in STN and ZI. **d** The normalized trial average of “go” responses for recorded cells from both STN and ZI. Each row is color-coded according to the region and the corresponding subtype of the cell. For the regions, blue and red indicate STN and ZI, respectively. For the subtypes, the ones with similar patterns shown in Fig. 2c-e (STN only) match the color, while the extra clusters shown in gray. **e** Single-cell trial-averaged traces from each cluster are displayed as colored thin lines, while the cluster-averaged traces are represented by black thick lines. The donut plots in the lower parts indicate the proportional distribution of STN and ZI cells in each cluster. **f** Single cells are projected on the 3D subspace of the “go” response, which are color-coded by their clusters (left) and the regions (right). **g** Histograms for the probability distribution of the cells from STN and ZI along each PC axis. **h**-**k** Same as **d**-**g** but under the “stop” condition. **l**-**o** Same as **d**-**g** but under the “rLick” condition. The red arrow highlights the differential distributions on the PC 2 between STN and ZI neurons.

To fairly compare the single-cell representation in the STN and the ZI, cells from the two regions are pooled for clustering analysis. The proportion of the cell numbers from each FOV is shown in Fig. 7c. Focusing on responses to locomotion and “reward-driven licking”, new clusters were identified beyond the ones shown in the clustering of STN neurons (Fig. 7d and e). In the “go” response, clusters showing *pre-move activation*, *long rising activity, or mild dipping* (last three clusters in Fig. 7e) are dominated by the ZI population. Meanwhile, the response subtypes that resembles “Go-2” and “Go-3”, which are mentioned, are dominated by STN neurons (Fig. 7e). To further analyze the discrepancy of response pattens beyond discrete clustering, we compared the distribution of single-cell responses in the feature subspace between the two regions (Fig. 7f and S7e). There is a slight bias showing in the axis of PC3 in the subspace of “go” response (Fig. 7g). Responses in the STN neurons show bias toward the positive direction of PC3, which represents the features of “Go-2” and “Go-3” subtypes; whereas the patterns of ZI neurons lean to the negative side, which represent the *long rising* feature (Fig. 7f and g). Apart from these nuances, the mean activity patterns at spontaneous locomotion (“go” and “stop”) are similar in the two regions, exhibiting large overlap in the feature subspace (Fig. 7d-k). However, the populational patterns of the STN and ZI are more distinguishable in the “rLick” responses. The clustering analysis identifies a cluster exhibiting “mild dropping” pattern, which was not clearly classified in the previous analysis focusing only in the STN (Fig. 7l-m). Additionally, it is worth to emphasize that the “rLick-1” and “rLick-4” subtypes contain relatively less ZI neurons (Fig. 7l-m). Importantly, the distribution of the neuronal responses in the subspace suggests that while responses in STN neurons are broadly covering the feature space, those in ZI neurons are more limited. The differences exhibit particularly on the axis of PC2, which is explained by the features of “rLick-1” and “rLick-4” subtypes (Fig. 7n, o, and S7g). The discrepancy in the distribution indicates that in response to reward-driven licking, neurons in the STN and the ZI share some common activity pattern, with the former expressing more diverse features than the latter. These results suggest that single-cell responses in the STN and ZI are to an extent distinguishable.

While the aforementioned description of behavioral representations focuses on the mean trace of activity, it lost the resolution in single trials. Indeed, we have observed that at the single-trial level, neuronal activities in the ZI are not as reliable as in that in the STN (Fig. S8). To compare the movement information encoded in both regions with the consideration of response reliability, we utilize linear regression to decode the moving speed and the licking patterns from neuronal single-trial activity. The results demonstrate that in both speed and licking intensity, while activity from ZI neurons can partially predict the movement, the performance is significantly higher when using STN activity (Fig 8a-d). Specifically, the prediction performance of the licking, in terms of coefficient correlation to the ground truth, is twice higher in the STN activity when compared with the ZI one (Fig. 8d). When looking into the predicted traces at the behavior transition, including “Go”, “Stop”, “rLick”, and “sLick”, predictions from STN neuronal activity outperform the ones from ZI. The difference is especially larger in the reward-driven licking (Fig. 8e), suggesting that ZI neurons encodes less, if any, information of the licking behavior. These indicates that while both regions encode locomotion and licking information, STN activity represents more faithfully than the ZI.

**Figure 8.**
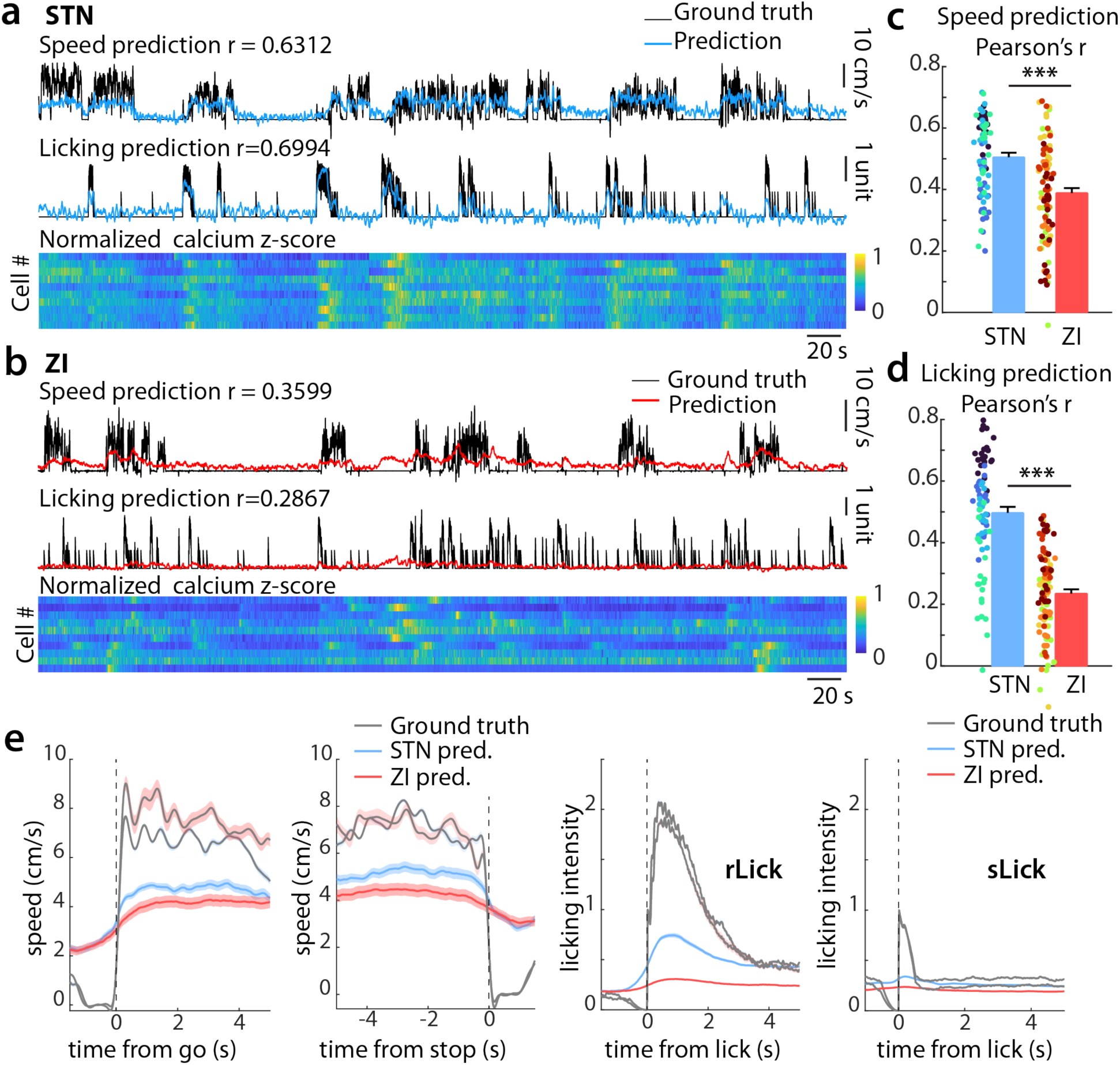
STN neurons encode more information in locomotion and licking than ZI neurons. **a, b** Examples of decoding locomotion speed and licking patterns from neurons in STN (**a**) and ZI (**b**). The heatmaps in the lower lanes are the normalized calcium responses of the corresponding sessions. **c**, **d** Quantification of the Pearson’s correlation coefficient between the predicted trace and the ground truth of speed (**c**) and licking (**d**). Dots with same color indicate the results from 20 replicates of a recording plane. *t*-test, *p<0.05, **p<0.01, ***p<0.001. **e** Predicted traces of speed (left two) and licking (right two) from STN and ZI neurons are aligned to “go”, “stop”, “rLick”, and “sLick”. The lines and the shades are respectively the mean and SEM of the average of events in each replicates (n=20*4 and 20*5 for STN and ZI, respectively).

## Discussion

### *In vivo* deep-brain imaging enables characterization of real-time neuronal activities and the cell identities

While the manipulation strategies allow for the investigation of the causality of neural activity and behavior, real-time recording captures more physiologically relevant correlations. Together with works in humans, monkeys, and rodents, the STN recordings are predominantly conducted using extracellular electrophysiology and reach the single-unit resolution through spike sorting. Although single-unit recording has provided substantial evidence for the functionality of the STN neurons, calcium imaging with cellular resolution offers a more straightforward identity of each cell. The cell identification provides precise spatial resolution, allowing for confident determination of signal positions within such a small structure for several days of recording. Moreover, the advantages of imaging in terms of cell identification surpass those of electrophysiology, particularly in the ease of integrating genetic tools and histology.

STN *in vivo* imaging is challenging due to its depth and small volume. The mouse STN is located around 4.5 mm beneath the brain surface, requiring a GRIN lens to penetrate this distance. The small, tilted, disc-like shaped STN (Fig. S1a and b) also presents challenges for accurate implantation. To address this, we utilize optically cleared thick slices to deterministically trace back the anatomical positions of the recorded cells (see Methods).

In this study, we established a surgical and imaging pipeline for monitoring the STN neuronal calcium activity while simultaneously tracking multiple behavioral parameters in real-time. This approach enables us to investigate the functional representation of single neurons within the STN. Through accumulating many recording sessions across days, we are allowed to capture the diverse calcium patterns under several behavioral and context conditions in individual neurons, which is required for investigating their various representations.

### The STN neurons are heterogeneous in their responses to a specific behavior

In line with single-unit recording studies, our results reveal functional subtypes of cells in the STN. Previous primate studies highlighted cells belonging to a subtype are tuned to a specific condition, such as “stop cells” or “switch cells” ^17,18^. In contrast, our findings demonstrate that a significant proportion of recorded neurons express distinct calcium dynamics under different behavioral conditions. Specifically, during self-initiated locomotion (“go” and “stop”) and non-reward licking (“sLick”), the calcium activities are relatively uniform but with subtle differences. The STN neurons are generally activated during both types of self-paced movement, aligning with other studies that observed movement-related activity ^14,15,25^. Interestingly, we further found that in response to reward-driven licking, four types of STN cells exhibit unique calcium activity patterns. We hypothesize that these distinctions may arise from the underlying cell type, upstream connectivity, or both. Notably, one of the subtypes (rLick-1) displays an inhibiting response, likely due to GABAergic GPe input that suppresses tonic STN firing. This type of cell tends to exhibit an inhibition prior to activation during locomotion onset (Go-2, Fig. 2c-e and 4f), which as well imply a stronger inhibitory input.

Moreover, this study obtained data from only four recording fields, which makes it challenging to draw conclusions about the spatial distribution of the functional subtypes. However, it is worth noticing that the diversity is observed within a single recording plane, which spans a width of less than 300 μm. This suggests that the variety of calcium responses originates from cells that are spatially intermingled, rather than being organized into distinct STN territories.

### Single STN neurons are capable of representing various behavioral conditions

Our results show that a majority of STN neurons respond to locomotion as well as various licking behaviors—both spontaneous and reward-driven—with distinct calcium activity patterns.

Most neurons show activation during internally driven movements, including locomotion and spontaneous licking. This aligns with the knowledge that the STN receives motor cortical inputs via the hyper-direct pathway along with pallidal inputs from the indirect pathway. It is well-established that striatal neurons, including those in direct and indirect pathways, are activated during movement ^27,28^. Consequently, cortical excitation combined with disinhibition from the indirect pathway leads to the activation of STN neurons. Furthermore, the STN has reciprocal connections to the mesencephalic locomotor region (MLR), which is known to regulate movements such as walking, grooming, rearing, and handling ^29,30^.

During reward-driven licking, on the other hand, these movement-activating neurons present various types of strong activity change, including inhibitory response and activations with distinct temporal patterns. Some behavioral events, which happen during reward-driven licking, are potentially represented by these activities.

First, continuous burst licking. While the cells that present fast-rise responses to the reward-driven licking (rLick-3 and rLick-4) mostly exhibit modest rising responses to the sparse, spontaneous licking, the patterns are not likely to be intuitively additive. It is shown in the analysis (Fig. 4) that, cells from the *prolonged activating* subtype “rLick-3” contain a higher percentage of *short activation* under spontaneous licking (sLick-2). Additionally, a proportion of the cells presenting activation to sparse licks show strong inhibition during reward-driven licking (rLick-1). These findings suggest that licking is encoded in a state- or context-dependent manner in single STN neurons, rather than through a simple additive process.

Second, action switching. While the reward-driven licking involves many trials with sudden stopping, the activity is distinct from that observed in non-reward, self-paced stopping in single neurons. This potentially suggests that the cue-directed stops (through the sound from the water port immediately prior to reward) are modulated differently from spontaneous stops. Such goal-directed stops are known to be influenced by the premotor cortico-STN connection, which facilitates rapid movement inhibition ^12,13^. Furthermore, our single-trial results reveal that the responses to reward-licking are similar regardless of whether stop events occur. We hypothesize that the STN neurons transmit neural signals, through both activation and inhibition, in response to reward cues not to encode deceleration *per se*, but to facilitate a transition between action states.

Finally, the dopamine-mediated reward prediction signal is also possibly modulated by the unique activities, especially the delayed type of activation. Indeed, reward amplitude has been shown to be encoded in rat STN ^23^. However, current evidence in our work is inadequate to determine the pure “reward” component in the reward-driven licking responses, which requires a more structured behavioral task.

Although it is challenging to decouple the influences of these concurrent events, it is clear that the state transition from self-initiated locomotion to reward-driven burst licking is encoded within the STN neuronal dynamics. Among the cells, some show less responsiveness to a specific behavior compared to others, yet most cells exhibit multiple behavioral representations. Additionally, whereas response clusters are somewhat interdependent across conditions, a large proportion of cells represent the behaviors in a mixed manner. In other words, the response pattern of a single neuron under one condition is inadequate to predict its pattern under others. The mixed multiple representations are recently described in the STN of monkeys, in which single units respond to both “go” trial and “cancel” trials in diverse ways ^31^. This suggests that the pathways for different motor programs overlap and converge in the STN neurons to some extent.

### Neural dynamical system and the circuitry implications

To address the functional implication of a population of neurons, the representational perspective is usually referred to, showing how single-cell activities are tuned by a specific condition. Meanwhile, in a dynamical model, the responses of individual neurons are expected to reflect hidden dynamical factors ^32^. In this work, the results demonstrate that from the aspect of representation, single STN neurons show behavior- and context-dependent representations, which obscure the establishment of a generalized underlying coding principle. Hence, the dynamical model is implemented.

Dynamical systems are commonly used in studies of cortical areas, particularly due to the availability of large-scale recordings. In this work, it is the first attempt to look into the populational dynamics in the STN. As the number of the cells recorded in a single field is limited, we took the trial-averaged calcium activities and pooled the data from all recording fields. In this manner, responses from different cells may not originate from the same recording trial, and as a result, only the average dynamics are captured. However, by reducing the dimensions from 80 cells to 3 PCs, we were able to clearly depict the population dynamics under each behavior, capturing most of the variance.

The neural trajectories in the dynamical model reveal that the activity during reward-driven licking, in contrast to other self-paced behaviors, extensively projects onto PC 2 and 3. This means the reward-related responses are captured by higher dimensions, or in other words, weighted combinations of single-cell activities, which are distinct from the neural states for locomotion. Furthermore, PC 1 and PC 2 in the system are found to be highly correlated with speed and licking intensity, respectively, indicating two major hidden factors in the population are capable of encoding these parameters. Critically, in the time windows of reward-driven licking, the neural trajectories follow the PC 1’s representation in speed—the higher speed at higher PC 1—prior to the licking onsets (Fig. 5e and 6c). However, upon the reward delivery and licking onset, the neural trajectory for the one with speed dropping does not decrease the PC 1 value, but travels along PC 2 and PC 3 instead (Fig. 6c). In summary, the system encodes the locomotion velocity and the licking intensity via two partially dependent hidden factors in a context-specific manner.

While the three principal components reflect the weighted average of the activity from the recorded neurons, hidden factors explaining specific motor behaviors or reward conditions imply that different sets of neurons in the STN are recruited in different behavioral programs. In addition, the functional representations of the hidden factors are context-dependent, indicating the same weighted combination of activities is involved in distinct functional pathways. Specifically, activities during cue-directed behaviors and spontaneous ones, which may be controlled by different neural pathways, are diverge from each other.

### Differential Neural Representations of Behavior in the ZI and STN

The ZI is a subthalamic region adjacent to the STN and is also an effective target for DBS in PD ^33^. It has been reported to be involved in physiological functions including sensory-motor integration, pain, and aversive behavior^34^. The ZI has a distinct cellular composition compared to the STN; for example, GABAergic neurons are wide-spread in the ZI but are rare in the STN ^33,35^. In addition, the ZI is reciprocally connected with a diverse range of regions, such as multiple cortical area, thalamus, and structures in midbrain and hindbrain^36^. Consequently, it is reasonable to expect that behavioral encoding in the ZI differs from that in the STN.

In our results, we identify functional discrepancies between populations in these two areas, both in mean activity and at single-trial level. For mean response patterns, although neurons in the STN and ZI share some features, the STN represents reward-driven licking by a greater diversity of response types compared to the ZI (Fig. 7n and o). At the single-trial level, we quantitatively observed that STN activity is more reliably associated with the selected behaviors than ZI activity. Furthermore, STN populations outperform ZI in the prediction in both locomotion and licking. Since the predictions were conducted using randomly selected, size-matched samples for model training and decoding, the results indicate that the information of the selected behaviors are more wide-spread in the STN population compared to the ZI.

These differences demonstrate that the two neighboring structures in the subthalamic region represent behaviors in distinguishable ways. This highlights the uniqueness of the encoding principle in the STN.

### Significance and perspectives

This study employs *in vivo* single-cell calcium imaging of the STN to establish correlations between neuronal activity and behavioral conditions. The results reveal functional heterogeneity and behavior-correlated neural dynamics within the STN. The diverse activities expressed by the STN neurons are shown to be context- and behavior-dependent, challenging the traditional perspective that primarily associates the STN with basic motor control. The observed functional diversity within the STN is spatially intermingled, which contradicts the conventional model of segregated functional domains within this structure. We further demonstrated the overlapping and convergency of inputs from the upstream areas. These results collectively explain the confounding off-target effects in clinical STN-DBS applications.

## Acknowledgments

The authors thank Fu-Chin Liu, Chiung-Chu Zoe Chen, Yi-Ping Hsueh, Shih-Chieh Lin, Ching-Lung Hsu, Hau-Jie Yau, and members of Wu laboratory for helpful discussions. We thank Ching-Lung Hsu, Shi-Bing Yang, Hau-Jie Yau, the IBMS AAV Core Facility (Institute of Biomedical Sciences, Academia Sinica) and the NTU AAV Core of National Taiwan University for providing AAVs used in this study. We thank the IMB Imaging Core Facility and IMB Animal Facility (Institute of Molecular Biology, Academia Sinica) for helpful supports. This work is supported by grants from the Academia Sinica (Y.-W. W.) and National Science and Technology Council (NSTC), Taiwan (Y.-W. W.; Brain Technology Grant: 112-2321-B-001-007; 113-2321-B-001-012), and Institute of Molecular Biology, Academia Sinica (Start-up fund, and SPP fund).

## Author contributions

Authors Contributions: B.S.W., Y.W.W., and M.Y.M. designed the experiments. B.S.W. performed the experiments. B.S.W. and Y.W.W. wrote the manuscript with contributions from M.Y.M.

## Figure legends

**Supplementary Figure 1.**
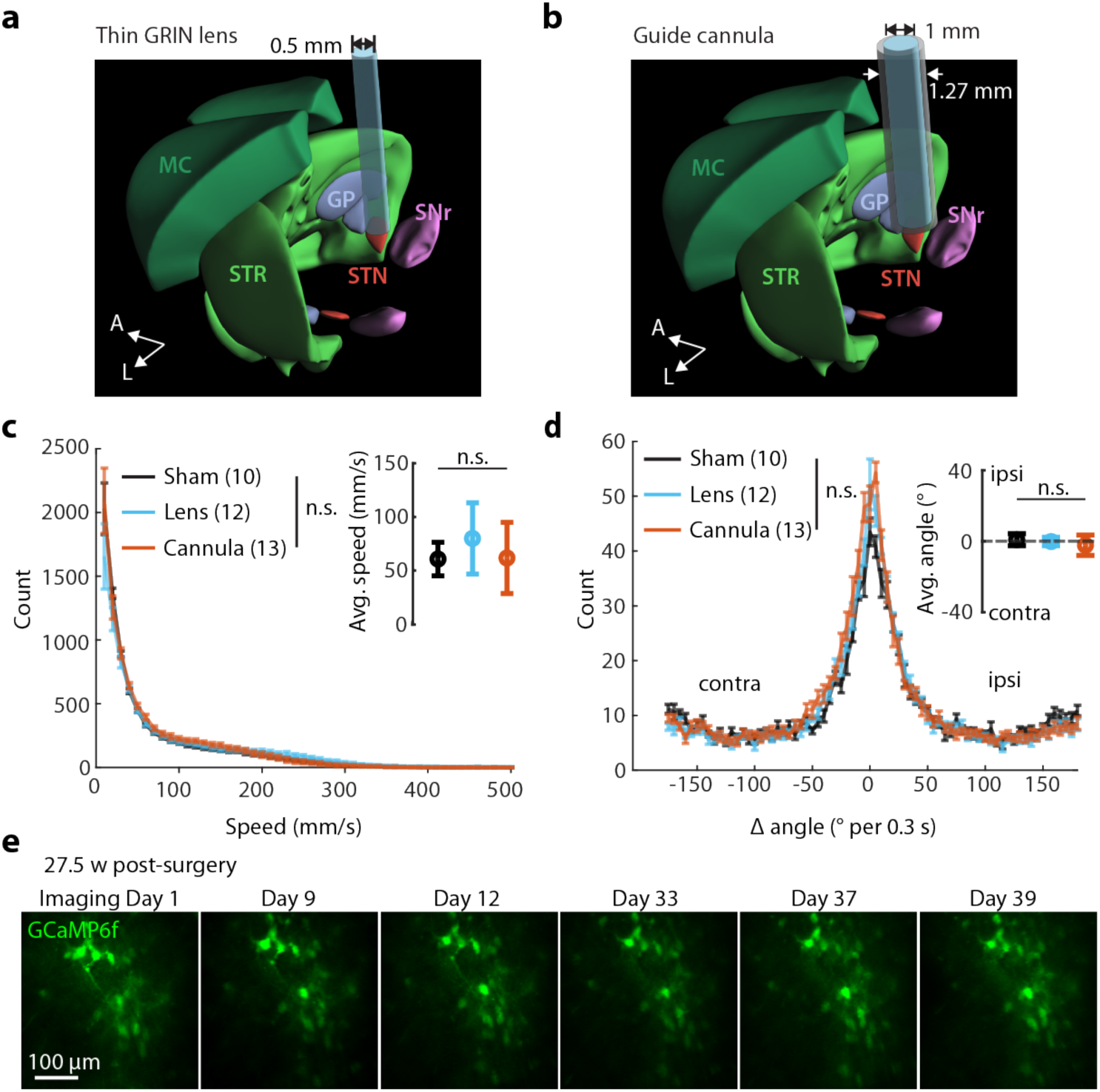
Stable deep-brain imaging minimally disrupts motor circuits. **a, b,** Schematics of the anatomical structures of cortico-basal ganglia regions and the implantation position of GRIN lens (**a**) and guide cannula (**b**). The implanted devices targeting subthalamic regions avoid damaging the cortico-basal ganglia circuitry. **c, d,** Mice with either GRIN lens or cannula implantation showed no significant difference in basic locomotion compared to sham control. **c,** Distribution of instantaneous speed (calculated per 0.033 ms) in open field test (5 minutes). *Inset,* average speed of 5 min. **d,** Distribution of rotational bias. *Inset*, average rotational bias in 5 min. Sham control, N = 10; GRIN Lens, N = 12; Cannula, N = 13. Two-way and one-way (inset) ANOVA. **e,** Representative images showing cross-day stability of imaging quality.

**Supplementary Figure 2.**
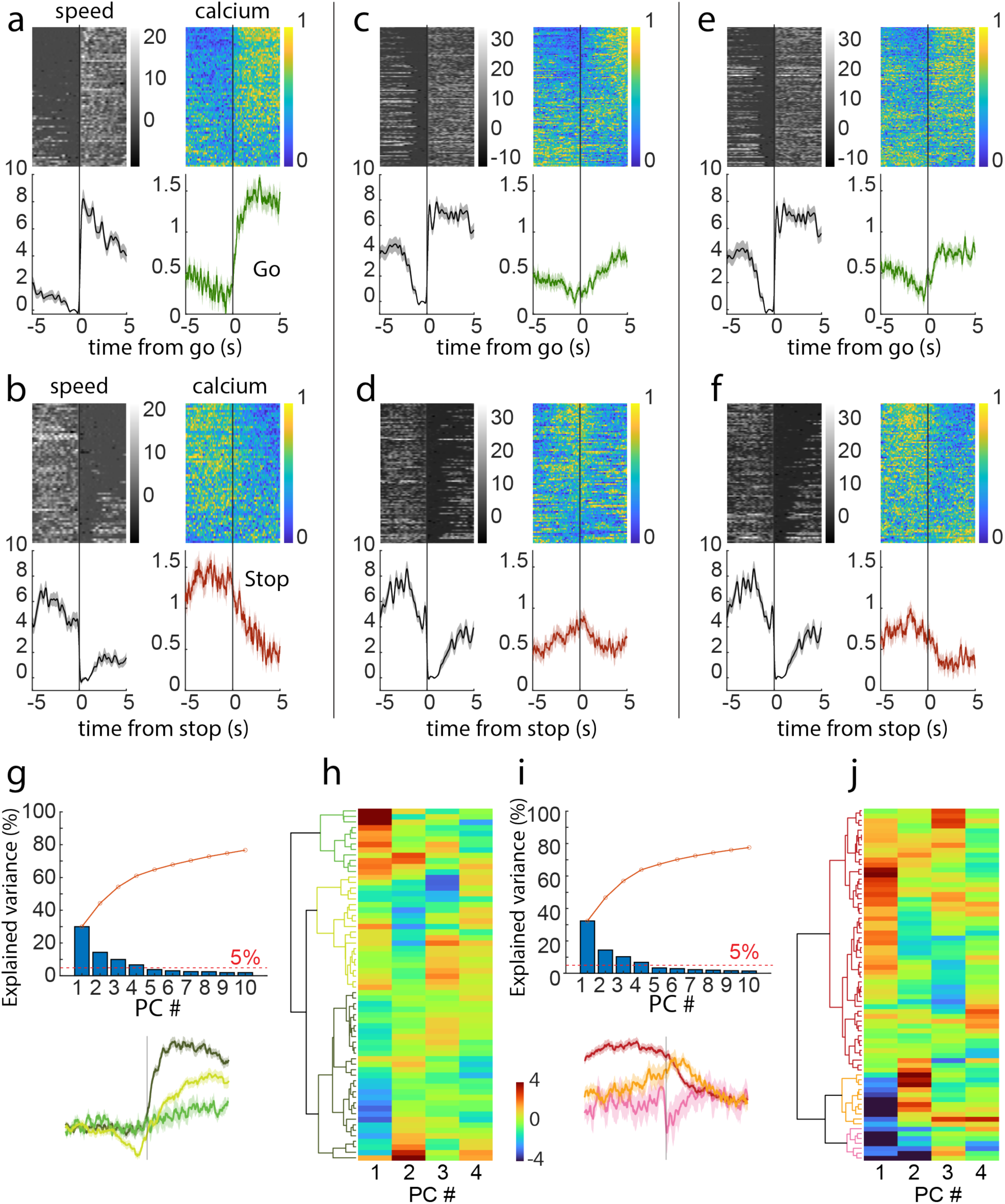
Examples of go and stop responses across single trials in single STN neurons and the clustering criteria. **a, b** Single-trial (upper panels) and mean (lower panels) traces of the speed and the calcium responses around “go” (a) and “stop” (b) from a single cell. **c, d** and **e, f** are traces of the other two example cells, respectively. **g** Explained variance of the first 10 PCs of “go” responses and the cumulated curve. **h** PCs explained more than 5% of variance (PC1∼4) are subjected to hierarchical clustering. The heatmap is the PC values of the “go” responses. **i**, **j** Same as **g**, **h** but for “stop” responses.

**Supplementary Figure 3.**
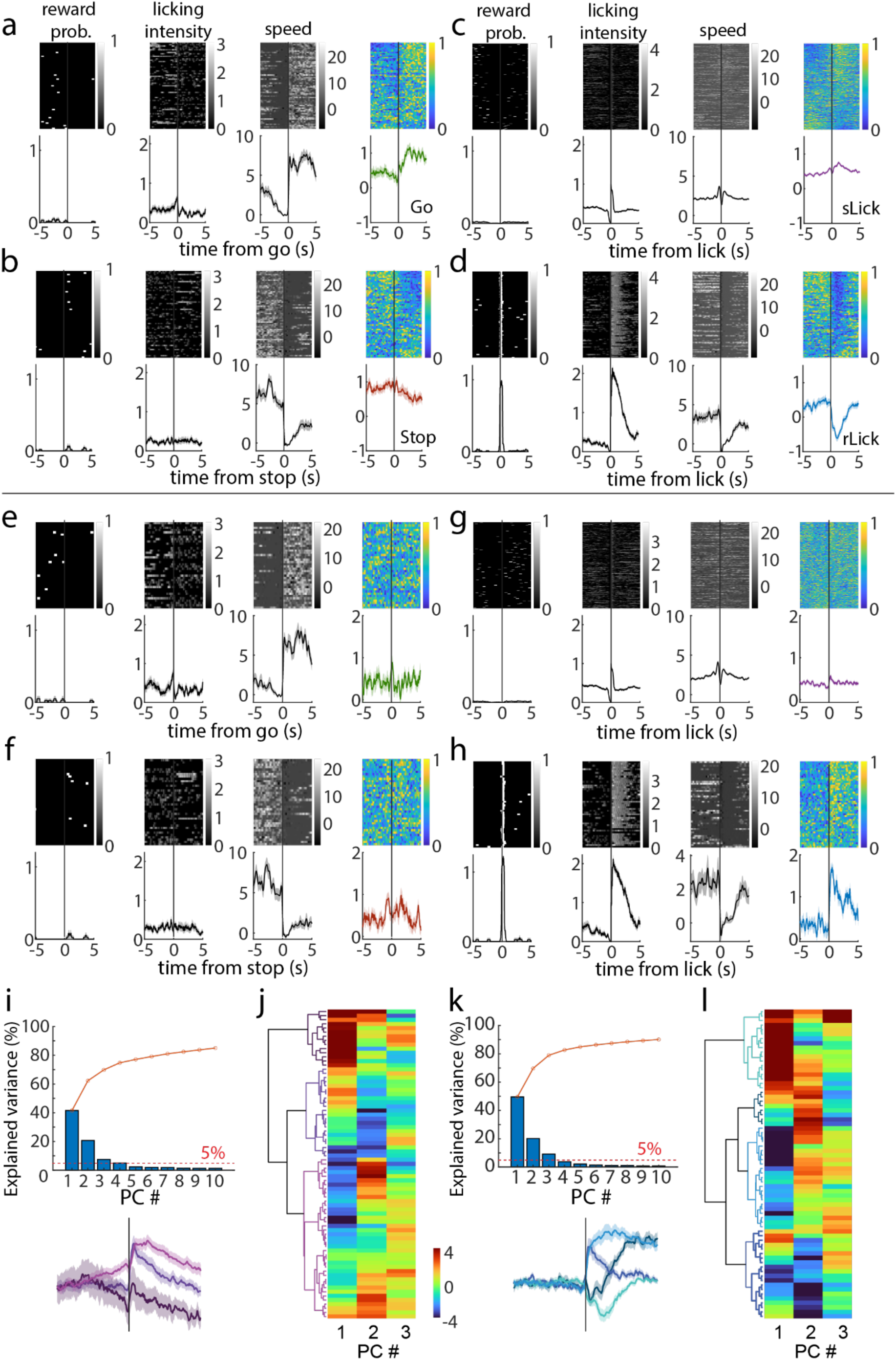
Single-cell, single-trial examples and the clustering criteria of sLick and rLick responses. **a-d** Single-trial and mean traces of reward probability, licking intensity, speed, and the calcium responses from a single cell under the four behavioral conditions. **e**-**h** Same as **a**-**d** but for another example cell. **i**, **j** Same as Fig. S2**g**-**h** but for “sLick”. **k**, **l** Same as Fig. S2**g**-**h** but for “rLick”.

**Supplementary Figure 4.**
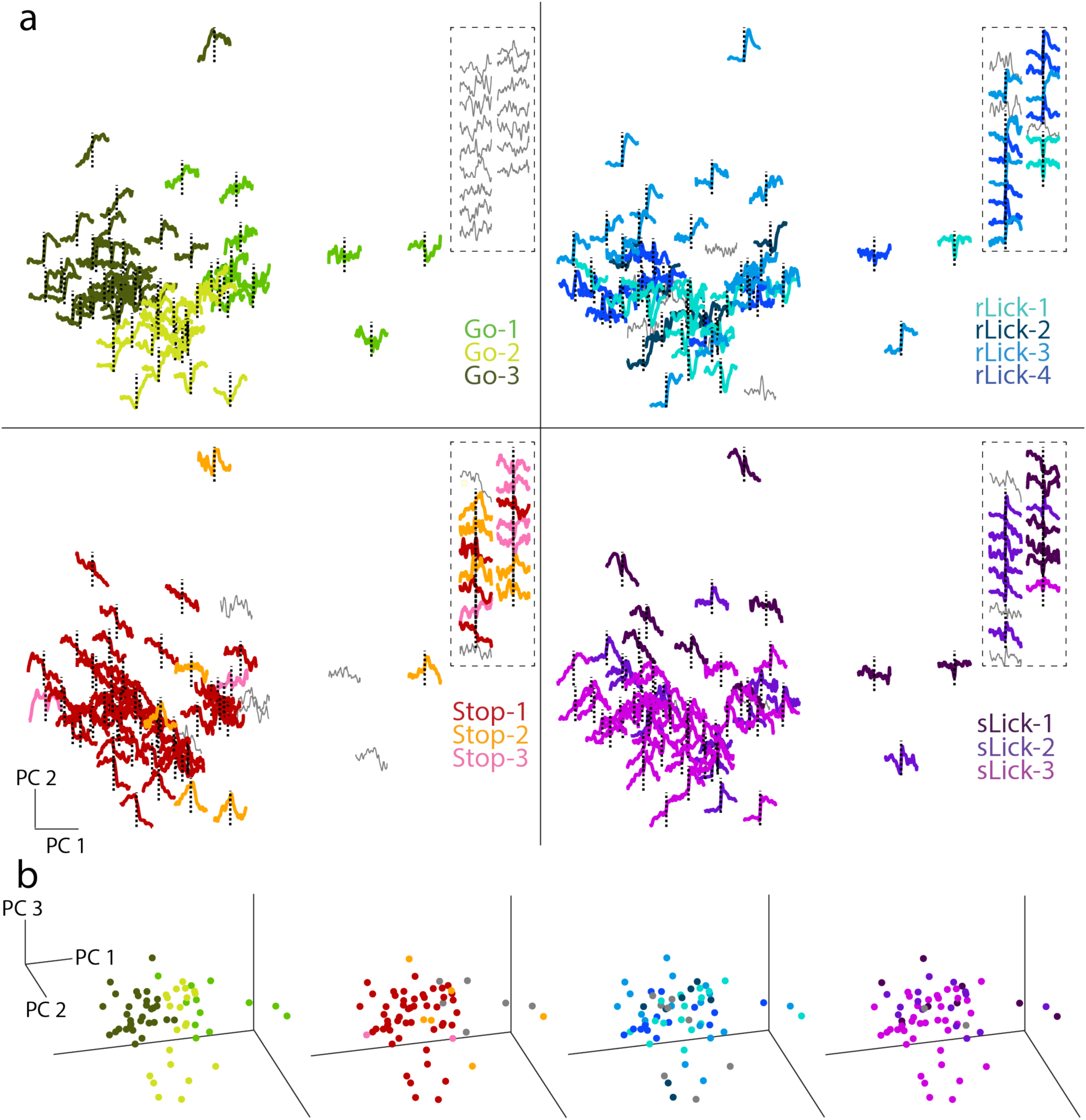
Cross-talking of single-cell calcium responses across behaviors in the “go” response subspace. Same as Fig. 4**a, b** but are mapped on the 2D and 3D subspace of “go” response.

**Supplementary Figure 5.**
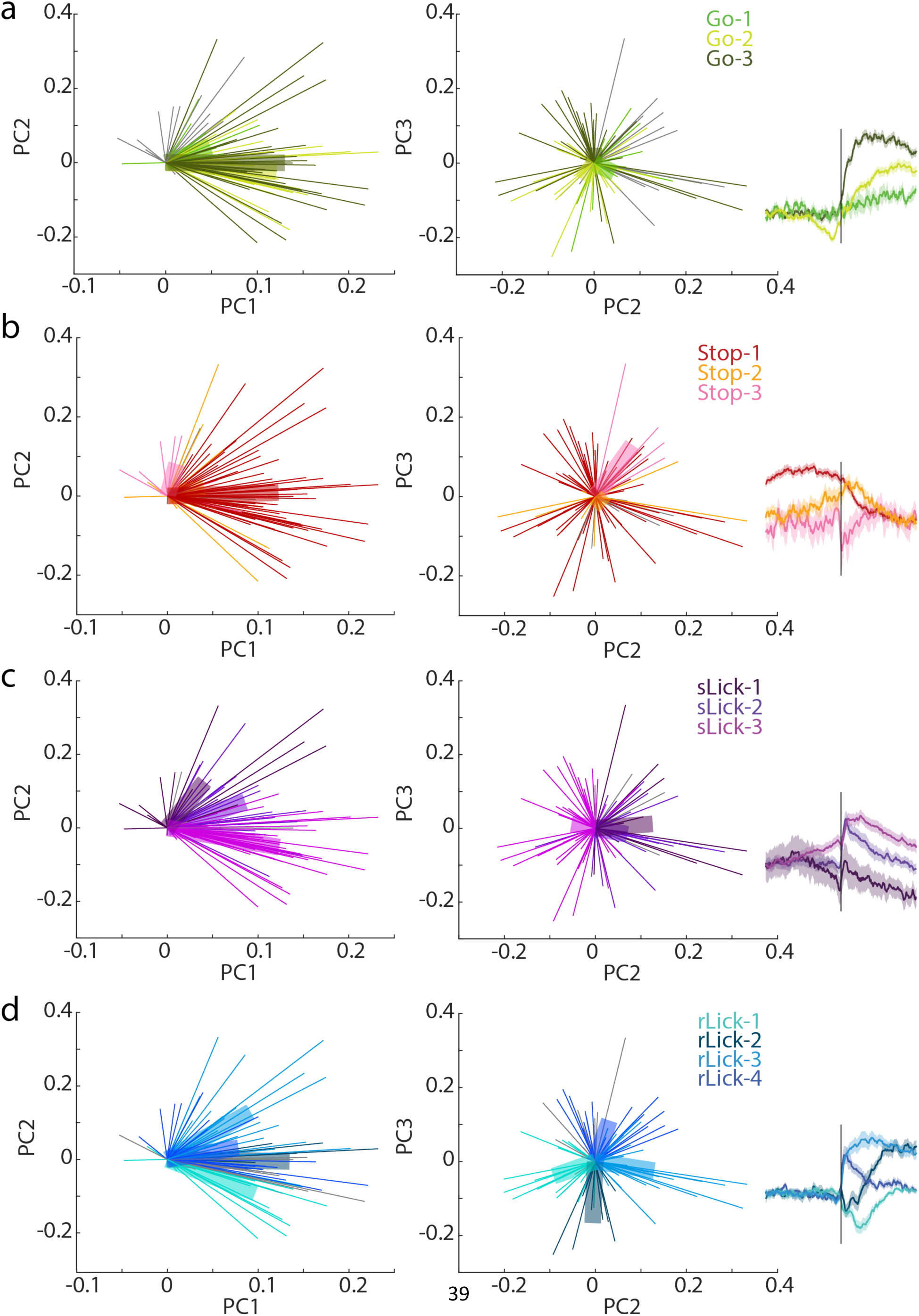
Single-cell contribution to the PC manifold. Thin lines are the coefficients (weights) vector of each cell’s responses for PC 1, 2, and 3. The vectors are color-coded by the responses subtypes and each thick bar is the average vector of each subtype.

**Supplementary Figure 6.**
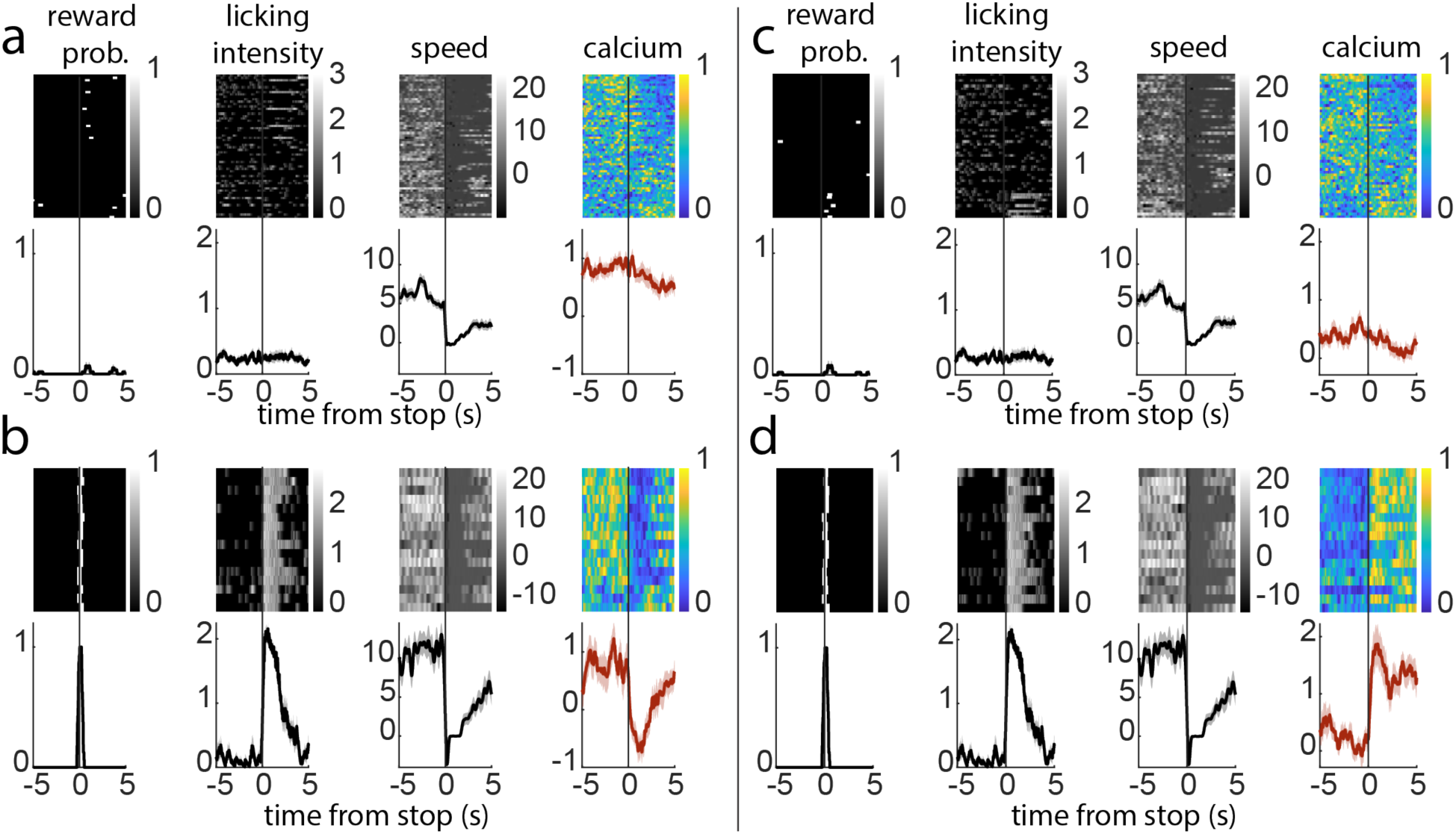
Single-trial traces demonstrating the divergence of responses to locomotion deceleration is dependent on reward-driven licking. **a** The single cell example of the responses to stops without reward (spontaneous stop, which is the condition defined as “stop” in the article). **b** Same as in (**a**) but under stops immediately followed by reward-driven licking. **c**, **d** Same as in **a**, **b** but for another example cell.

**Supplementary Figure 7.**
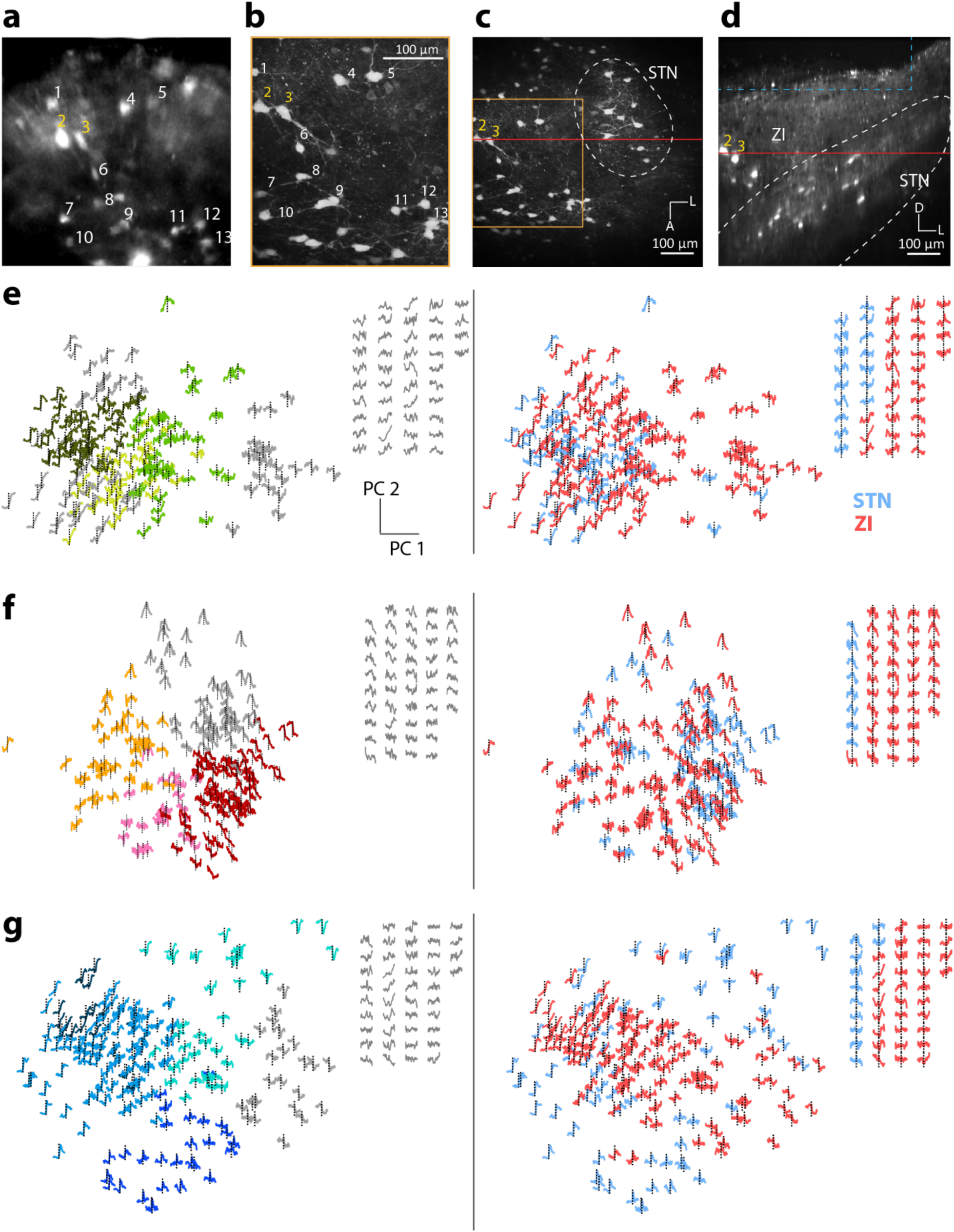
The *post hoc* validation in ZI and the 2D PC subspace of STN and ZI calcium responses across behavioral condition. **a-d** Representative images demonstrating the *post hoc* validation of recorded cells in the ZI. **a** Averaged image for the in vivo recording field. **b** Cells in the fixed slice that match those in **a**. **c** Zoom out view of **b**. **d** Orthogonal view of **c**. **e-g** Mean traces of single cells from both STN and ZI mapped on the 2D subspace of “go” (**e**), “stop” (**f**), and “rLick” (**g**). Traces are color-coded by the response subtypes (left panels) and the regions (right panels).

**Supplementary Figure 8.**
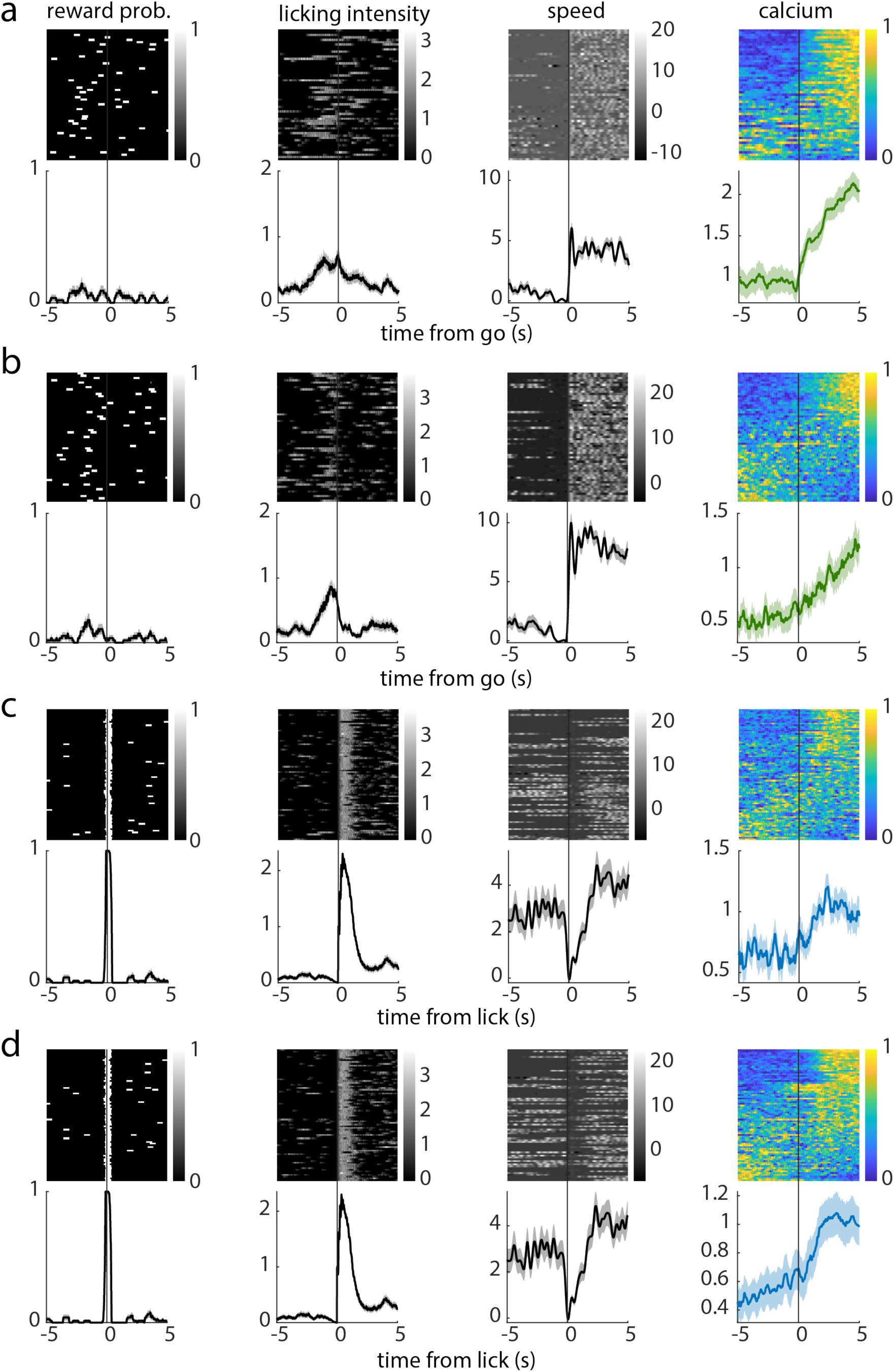
Single trial examples showing relative unreliable responses of ZI neurons under “go” and “rLick”. **a, b** Single-cell, single-trial traces of behaviors and calcium responses under “go”. **c**, **d** Same as a, b but under “rLick.

## STAR Methods

### Key resources table

#### Experimental model

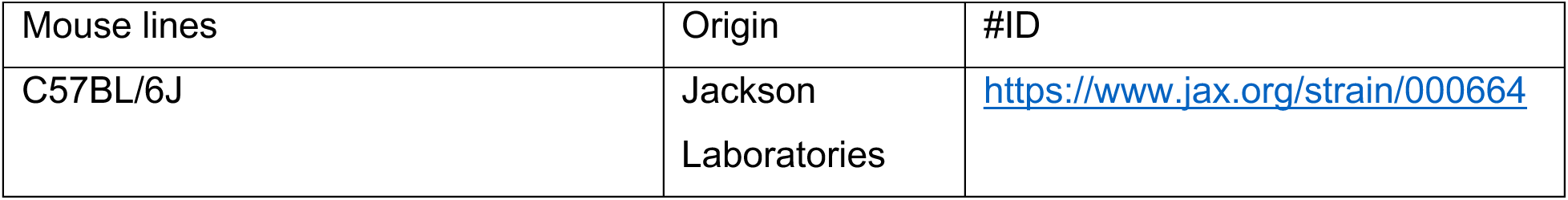

#### Software and Algorithms

|  |  |
| --- | --- |
| ImageJ v1.54f | NIH |
| MATLAB | Mathworks, USA |
| Prism v10.0.0 | Graphpad |
| Imaris |  |
| ZEN blue/black (SIM deconvolution) | ZEISS |

### RESOURCE AVAILABILITY

#### Lead contact

Dr. Yu-Wei Wu

Institute of Molecular Biology, Academia Sinica, Taipei 115, Taiwan; +886-2-2789-9334 (Tel)

Mr. Bing-Shiuan Wu

Institute of Molecular Biology, Academia Sinica, Taipei 115, Taiwan; +886-2-2789-9334 (Tel)

#### Material availability statement

All materials generated in this study are available from the corresponding author upon reasonable request. Any unique reagents, data, or protocols developed for this study will be made available under standard material transfer agreements (MTAs). Publicly available datasets used in this study can be accessed as indicated in the Methods section.

#### Data and code availability

The datasets generated and analyzed during the current study are available from the corresponding author upon reasonable request. All code used for data analysis is also available upon request and will be shared under a standard open-source license. Publicly available datasets referenced in this study can be accessed as described in the Methods section. Any additional information required to reproduce the study’s findings is available from the corresponding author upon reasonable request

### EXPERIMENTAL MODEL AND SUBJECT DETAILS

#### Mouse lines

Male C57BL/6J mice (8-12 weeks old) were used for the open field test. Mice were housed in groups of 4-5 per cage in a temperature-controlled environment (22 ± 1°C) with a 12-hour light/dark cycle. 3 weeks before behavioral assay, the mice were transfer to a room with reversed light/dark cycle. Food and water were available ad libitum. All procedures were conducted in accordance with the institutional guidelines and approved by the Institutional Animal Care and Use Committee (IACUC) at Academia Sinica.

#### Stereotaxic surgeries

Adult mice over two months are used for the surgeries. The mice were anesthetized by isoflurane (1 – 5%) and stabilized in a stereotaxic frame before the surgeries.

##### Viral injection

For the GCaMP6f expression in the STN, a craniotomy was done above the target prior to the viral injection. The viral vectors AAV9-CAG-GCaMP6f-WPRE-SV40 (Addgene-100835; qTiter: 2.34*1013 GC/mL) were 10 times diluted with saline. 0.5 μL of the solution was microinjected to either side of the STN (AP -1.8 mm, ML ±1.67 mm, DV -4.5 mm) at a rate of 1 nL/sec through NanoJect 3 (Drummond Scientific, USA).

For the transsynaptic tracing, the viral vectors AAV9-hSyn-Cre-P2A-dTomato (Addgene-107738; qTiter: 2.7*1013 GC/mL) and AAV9-Ef1a-Flpo (Addgene-55637; qTiter: 1.6*1013 GC/mL) are injected in pair into two cortical regions, either M1 (rostral: AP -1.8 mm, ML ±1.67 mm, DV -4.5 mm; caudal: AP -1.8 mm, ML ±1.67 mm, DV -4.5 mm) paired with M2 (rostral: AP +2 mm, ML ±0.5 mm, DV -0.8 mm or AP +2.5 mm, ML ±1 mm, DV -0.8 mm; caudal: AP +1 or +1.5 mm, ML ±1.5 mm, DV -0.8 mm), or M1 paired with S1 (rostral: AP +0.5 mm, ML ±2.5 mm, DV -0.95 mm; caudal: AP -0.5 mm, ML ±2 mm, DV -0.8 mm). Both rostral and caudal sites are injected with 50 nL. For the recombinase-dependent reporters, AAV9-Ef1a-DIO-EYFP (Addgene-27056; qTiter: 8.7*1013 GC/mL) and AAV9-Ef1a-fDIO-tdTomato (Addgene-128434; qTiter: 8.1*1013 GC/mL) were 10 times diluted with saline and then 1:1 mixed. 100 to 200 nL of the mixed solution was injected into the STN. The samples were collected 4 weeks after the viral injection.

##### Implantation of GRIN lens

The GRIN lens is access to the STN by loading into an implanted guided cannula or directly implanting to the target. For guided cannula implantation, the stainless cannula with a 1.3 mm diameter was attached to the polished cover glass, which allowed imaging through a 1 mm-width GRIN lens (Inscopix, PN 130-000143, NA 0.5). The tissue above the target is removed via aspiration along with flushing mammalian Ringer’s solution. The cannula is lowered to the position above the STN (DV, -4.3 mm). For the directly-implanted GRIN lens, following creating a track via a 27-gauge needle, a 0.5 mm-width GRIN lens (GRINTECH, NEM-050-50-00-920-S-1.5p, NA 0.5) is implanted to the position above the STN (DV, -4.4 mm). At the end of the surgery, a custom-made head plate was secured to the skull using dental cement, enabling the head-fix imaging sessions. Over a month was allowed for recovery and viral expression.

#### Unstructured behavior assay

##### Experimental setting

Before the imaging session, the mice had undergone water restriction (1 mL per day) and habituated to the head-fix behavior stage for at least two days. During the imaging, the mouse was head-fixed and allowed to run on a running wheel. Approximately 5 μL droplets of 4% sucrose solution were delivered at random timing through the blunt needle connected with a water port.

##### Behavioral data acquisition

During the recording session, the rotation of the running wheel was recorded by the established rotary encoder. The reward-delivery timing and the encoder data are synchronized to the imaging. Body part movements were captured by an infrared camera (BFS-U3-13Y3, FLIR, US), which is triggered upon the two-photon recording at 50 frames-per-second (fps).

##### Behavioral data analysis

The data are resampled to 50 Hz to match the frame rate of the camera. After the speed was converted from the encoder data, a threshold (1.7 cm/sec) was set to define the moving episodes. The onset and the offset of the moving episodes lasting longer than 4 seconds are described as “go” and “stop”, while the onset for the moving less than 4 seconds is defined as “short run”. The licking events were captured by the detection of tongue movement in the video through DeepLabCut (Nath et al., 2019). After converting the licking events into binary data, a kernel with a 0.5-second linear decay was applied along the binary series. The starting points of all discrete licking bouts post-convolution are defined as licking onsets. For separating the reward-driven licking from spontaneous licking conditions, as well as separating reward-driven stops from spontaneous ones, the reward probability traces in the window of ±3 sec from the transition points are subjected to correlation analysis. There were clear-cut difference in the paired correlation between “with reward” and “without reward” trials and can be simply separated by hierarchical clustering.

#### Two-photon imaging

##### In vivo deep-brain imaging

A two-photon microscopy system (FemtoSmart Dual, Femtonics, Hungary) with a 20x//1.0 NA water immersion objective lens (XLUMPLFLN Objective, Olympus) was used in both *in vivo* and slice imaging. 920 nm excitation laser (Chameleon Ultra II, Coherent, USA) combined with a 490-550 nm bandpass filter was used in resonant scanning mode to detect *in vivo* calcium activity. The videos were recorded under a resolution of 512 x 512 and a frame rate of 31 Hz. Each recording session lasted for 5 minutes. 2 to 4 sessions of imaging were acquired within a single day, with each mouse being recorded for 3 to 4 non-consecutive days.

##### Thick slice imaging for post hoc implant site verification

The brains were collected after the mice were perfused with 4% PFA. 1 mm-thick slices containing the STN were sectioned by vibratome. For post hoc validation, the brains were horizontally sectioned from Bregma DV -4.0 mm to -5.0 mm. For transsynaptic tracing, 1 mm-thick sagittal slices were collected. The thick brain slices were then cleared following the ScaleSQ protocol (Hama et al., 2015). After tissue clearance, brain sections were mounted in the ScaleS4 solution and ready for two-photon imaging.

For defining the STN boundary, 820 nm excitation laser (Chameleon Ultra II, Coherent, USA) combined with a 450-490 nm bandpass filter was used in the galvo scanning mode to image the GCaMP6f-expressing cells as well as the brain area structures via auto-fluorescence. Image stacks were acquired with 1024 x 1024 pixels and 0.68 µm/pixel resolution. The zoomed-out view was stitched from tile-scanned images with the Stitching plugin in ImageJ (NIH, USA).

### QUANTIFICATION AND STATISTICAL ANALYSIS

#### Extraction of calcium dynamics and image preprocessing

The calcium imaging data are output from the MESc data acquisition software (Femtonics Ltd, Budapest, Hungary). The motion correction and the segmentation are done by Suite2P (Pachitariu *et al.*, 2017). The calcium traces are z-scored with the offset term being the baseline, which is defined by the mode of the signals. The traces are smoothed and resampled to 50 Hz to match the camera frame rate.

#### Calcium response significance definition

After extraction of the peri-event activities with a 10-second time window, the single-trial traces from each cell underwent a statistical test to determine the significance of the response. The mean values from the first bin (1.4-sec bin size) of single-trial traces, which is assumed to be the baseline of the response, are compared to those in the maximum bin and the minimum bin separately by paired student t-test. The cell with either the maximum or minimum bin significantly different (p<0.05) from the first bin is defined to be responsive. The non-responsive cells in a specific behavioral condition are excluded from the clustering analysis.

#### Cross correlation between calcium activity and behavior variables

Normalized weighted cross-correlations were calculated between the calcium trace and the corresponding behavior trace (either speed or licking) in each single trial through MATLAB functions. The time lags refer to the shifting of the calcium traces.

#### Calcium response clustering

The clustering was based on the trial averages of each cell under each behavioral condition. The response traces were subjected to principal component analysis (PCA) in the dimension of time. The first four principal components (PCs) of the response traces are taken in the “go” and “stop” conditions, while three PCs are considered in the two types of licking. The differences come from the percentage of the explained variance in each PC.

The values of the PCs in all the responsive cells are subjected to hierarchical clustering using inner squared distance (minimum variance algorithm) for the linkage method in the MATLAB function. Clustering cutoff for each behavioral conditions: “go”, 12; “stop”, 13; “rLick”,12; “sLick”,15.

#### Population dynamics

The mean response traces under the six mentioned behavioral conditions are concatenated in the temporal dimension, which means each cell would have six segments of 10-second-long trace in a row. The 80 cells with the concatenated traces are subjected to PCA in the dimension of cells (80 dimensions). The first three PCs are selected to be the axes of the three-dimensional subspace. The displayed trajectories were smoothed by the Savitzky-Golay filter for 50 data points (∼1 sec).

#### Behavior decoding

The decoding was performed with the function “fitrlinear” in MATLAB. The regression models were trained with least-squares algorithm under Lasso regularization. Data from each sample was undergone 20 replications of model training and prediction, with the involved neurons and trials are randomly selected and match the data size in every iteration.

